# Boosting with Omicron-matched or historical mRNA vaccines increases neutralizing antibody responses and protection against B.1.1.529 infection in mice

**DOI:** 10.1101/2022.02.07.479419

**Authors:** Baoling Ying, Suzanne M. Scheaffer, Bradley Whitener, Chieh-Yu Liang, Oleksandr Dmytrenko, Samantha Mackin, Kai Wu, Diana Lee, Laura E. Avena, Zhenlu Chong, James Brett Case, LingZhi Ma, Thu Kim, Caralyn Sein, Angela Woods, Daniela Montes Berrueta, Andrea Carfi, Sayda M. Elbashir, Darin K. Edwards, Larissa B. Thackray, Michael S. Diamond

## Abstract

The B.1.1.529 Omicron variant jeopardizes vaccines designed with early pandemic spike antigens. Here, we evaluated in mice the protective activity of the Moderna mRNA-1273 vaccine against B.1.1.529 before or after boosting with preclinical mRNA-1273 or mRNA-1273.529, an Omicron-matched vaccine. Whereas two doses of mRNA-1273 vaccine induced high levels of serum neutralizing antibodies against historical WA1/2020 strains, levels were lower against B.1.1.529 and associated with infection and inflammation in the lung. A primary vaccination series with mRNA-1273.529 potently neutralized B.1.1.529 but showed limited inhibition of historical or other SARS-CoV-2 variants. However, boosting with mRNA-1273 or mRNA-1273.529 vaccines increased serum neutralizing titers and protection against B.1.1.529 infection. Nonetheless, the levels of inhibitory antibodies were higher, and viral burden and cytokines in the lung were slightly lower in mice given the Omicron-matched mRNA booster. Thus, in mice, boosting with mRNA-1273 or mRNA-1273.529 enhances protection against B.1.1.529 infection with limited differences in efficacy measured.

## INTRODUCTION

Since the inception of the SARS-CoV-2 pandemic in late 2019, almost 400 million infections and 5.8 million deaths have been recorded (https://covid19.who.int). Several highly effective vaccines targeting the SARS-CoV-2 spike protein encompassing multiple platforms ((lipid nanoparticle encapsulated mRNA, inactivated virion, or viral-vectored vaccine platforms (Graham, 2020)) were developed and deployed rapidly with billions of doses administered (https://covid19.who.int). These vaccines use the SARS-CoV-2 spike protein from historical strains that circulated during the early phases of the pandemic in 2020 and have reduced the numbers of infections, hospitalizations, and COVID-19-related deaths. Despite the success of COVID-19 vaccines, the continued evolution of more transmissible SARS-CoV-2 variants with amino acid substitutions, deletions, and insertions in the spike protein jeopardizes the efficacy of global vaccination campaigns (Krause et al., 2021).

The SARS-CoV-2 spike protein engages angiotensin-converting enzyme 2 (ACE2) on the surface of human cells to facilitate entry and infection (Letko et al., 2020). The S1 fragment of the spike protein contains the N-terminal (NTD) and receptor binding (RBD) domains, which are targets of neutralizing monoclonal (Barnes et al., 2020; Cao et al., 2020; Pinto et al., 2020; Tortorici et al., 2020; Zost et al., 2020) and polyclonal antibodies (Rathe et al., 2020). In late November of 2021, the Omicron (B.1.1.529) variant emerged, which has the largest number (>30) of amino acid substitutions, deletions, or insertions in the spike protein described to date. These changes in the spike raise concerns for escape from protection by existing vaccines that target early pandemic spike proteins. Indeed, reduced serum neutralization of B.1.1.529 virus (Edara et al., 2021; Pajon et al., 2022) and large numbers of symptomatic breakthrough infections with B.1.1.529 have been reported in vaccinated individuals (Buchan et al., 2022; Christensen et al., 2022; Elliott et al., 2021).

Here, we evaluated the antibody responses and protective activity against B.1.1.529 Omicron virus of a preclinical version of the current Moderna vaccine, mRNA-1273, or an Omicron-targeted vaccine, mRNA-1273.529, designed with sequences from the historical Wuhan-1 or B. 1.1.529 spike genes, respectively, in the context of a primary (two-dose) immunization series or third-dose boosters. We hoped to define serum antibody correlates of protection against B. 1.1.529, determine the likelihood and significance of breakthrough infections, define the differences in immunogenicity and protection of homologous and heterologous mRNA vaccine boosters, and evaluate the activity of mRNA-1273.529 in the context of a primary immunization series against historical and variants of concern strains that emerged prior to Omicron.

After a primary immunization series, mRNA-1273.529 induced antibody responses that efficiently neutralized viruses displaying B.1.1.529 spike proteins but poorly inhibited infection of viruses expressing spike proteins of a historical strain or key variants (*e.g*., Beta (B.1.351) and Delta (B. 1.617.2)). Whereas a primary immunization series with a high-dose mRNA-1273 formulation conferred protection against both historical and B.1.1.529 viruses, a low-dose series, which induced levels of neutralizing antibodies against WA1/2020 strains that correspond to those measured in human serum (Wu et al., 2021), protected against WA1/2020 but did not control viral infection or inflammation in the lungs of B.1.1.529-challenged mice. Boosting with either mRNA-1273 or Omicron-matched mRNA-1273.529 vaccine increased levels of neutralizing antibodies and protection against B.1.1.529, although higher levels of neutralizing antibodies and lower levels of lung infection and inflammation were observed in mice boosted with mRNA-1273.529. Thus, the levels of vaccine-induced immunity that protect against historical or other variant SARS-CoV-2 strains (Ying et al., 2021) fail to prevent breakthrough infection by B.1.1.529 virus, necessitating boosting with either matched or unmatched vaccines.

## RESULTS

### Antibody responses against B.1.1.529 in K18-hACE2 mice

We evaluated the antibody response against B.1.1.529 after immunization with a preclinical version of mRNA-1273 that encodes for the prefusion-stabilized spike protein of SARS-CoV-2 Wuhan-1 strain (Corbett et al., 2020). We used K18-hACE2 transgenic mice, which are susceptible to severe infection after intranasal inoculation by many SARS-CoV-2 strains (Chen et al., 2021b; Winkler et al., 2020). Groups of 7-week-old female K18-hACE2 mice were immunized twice over three weeks by intramuscular route with 5 or 0.1 μg doses of mRNA-1273 or a control mRNA vaccine (**Fig 1A**). A lower vaccine dose arm was included for evaluating correlates of protection, as we expected breakthrough infections in this group. Serum samples were collected three weeks after the second dose, and IgG responses against spike proteins (Wuhan-1 and B.1.1.529) were evaluated by ELISA. We confirmed that equivalent amounts of antigenically intact Wuhan-1 and B.1.1.529 spike (proline-stabilized, S2P) and RBD proteins were adsorbed based on detection with sarbecovirus cross-reactive monoclonal antibodies (VanBlargan et al., 2021) (**Fig 1B**). Antibody responses against both the Wuhan-1 and B.1.1.529 spike proteins were robust after two immunizations with mRNA-1273. For the 5 μg dose, mean serum endpoint titers ranged from ~4,000,000 to 800,000 against the Wuhan-1 and B.1.1.529 spike proteins and 1,000,000 and 40,000 for the Wuhan-1 and B.1.1.529 RBD, respectively (**Fig 1C-D**). For the 0.1 μg dose, approximately 10-fold lower serum IgG responses against the spike and RBD proteins were measured (**Fig 1E-F**). The most noticeable difference in IgG responses after mRNA-1273 vaccination was the reduced titer against the B.1.1.529 RBD compared to Wuhan-1 RBD protein (**Fig 1D and F**).

**Figure 1.**
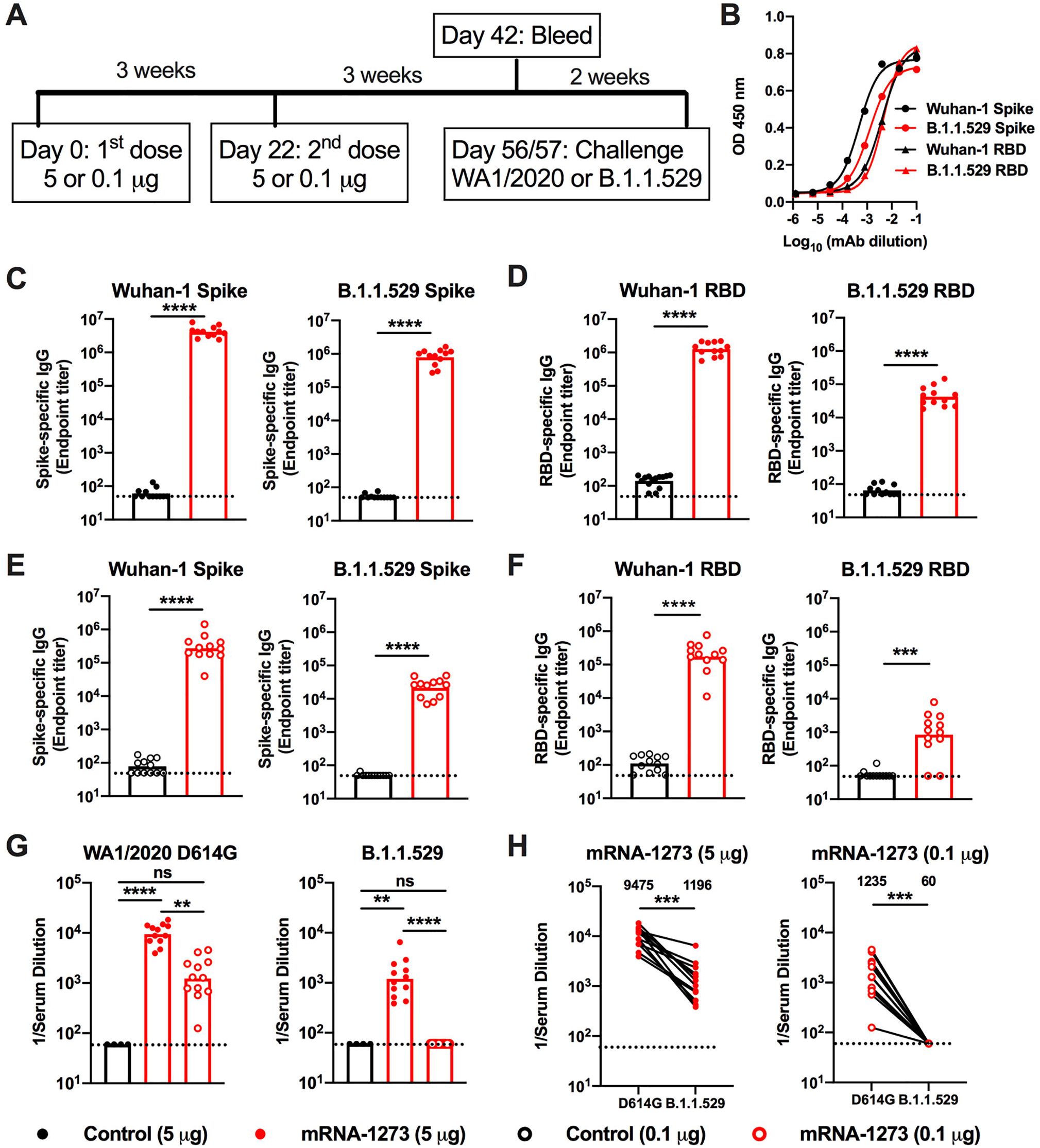
Antibody responses of mRNA vaccines in K18-hACE2 mice. Seven-week-old female K18-hACE2 transgenic mice were immunized with 5 or 0.1 μg of mRNA vaccines. **A**. Scheme of immunizations, blood draw, and virus challenge. **B**. Binding of sarbecovirus cross-reactive monoclonal antibodies to Wuhan-1 and B.1.1.529 spike and RBD proteins. **C-F**. Serum IgG responses at three weeks after the second 5 μg (**C-D**) or 0.1 μg (**E-F**) dose of mRNA vaccines (control or mRNA-1273) against indicated spike (**C, E**) or RBD (**D, F**) proteins (n = 12, two experiments, boxes illustrate geometric mean values, dotted lines show the limit of detection (LOD)). **G**. Serum neutralizing antibody responses three weeks after second vaccine dose as judged by focus reduction neutralization test (FRNT) with WA1/2020 D614G (*left panel*) and B.1.1.529 (*right panel*) in mice immunized with 5 or 0.1 μg of control (n = 4) or mRNA-1273 (n = 12) vaccines (two experiments, boxes illustrate geometric mean titer (GMT) values, dotted lines show the LOD). **H**. Paired analysis of serum neutralizing titers against WA1/2020 D614G and B.1.1.529 from individual mice (data from **G**) from samples obtained three weeks after the second 5 μg (*left panel*) or 0.1 μg (*right panel*) dose of mRNA-1273 (n = 12, two experiments, dotted lines show the LOD). GMT values are indicated at the top of the graphs. Statistical analyses. **C-F**: Mann-Whitney test; **G**: One-way ANOVA with Dunn’s post-test; **H**: Wilcoxon signed-rank test (** *P* < 0.01; *** *P* < 0.001; **** *P* < 0.0001).

We characterized functional antibody responses by measuring the inhibitory effects of serum on SARS-CoV-2 infectivity using a focus-reduction neutralization test (FRNT) (Case et al., 2020b) and fully-infectious SARS-CoV-2 WA1/2020 D614G and B.1.1.529 strains (**Fig 1G-H and S1**). Due to the limited amount of sera recovered from live animals, we started dilutions at 1/60, which is just above the estimated level of neutralizing antibodies associated with protection in humans (Khoury et al., 2021). For the 5 μg dose, the mRNA-1273 vaccine induced robust serum neutralizing antibody responses against both WA1/2020 D614G and B.1.1.529 (**Fig 1G-H and S1**). However, the geometric mean titers (GMTs) of neutralization were ~8-fold lower (*P* < 0.001) against B.1.1.529, which agrees with data from human antibodies (Cameroni et al., 2021; Cao et al., 2021; Cele et al., 2021; Dejnirattisai et al., 2022; VanBlargan et al., 2022; Wilhelm et al., 2021). For the 0.1 μg mRNA-1273 vaccine dose, we observed ~8-fold less (*P* < 0.01) serum neutralizing activity against WA1/2020 D614G compared to the higher vaccine dose. Serum from mRNA-1273-vaccinated mice with the 0.1 μg dose also showed large (>20-fold, *P* < 0.001) reductions in neutralization of B.1.1.529, with all values assigned to the 1/60 limit of detection (**Fig 1H**).

### Protection against B.1.1.529 by mRNA-1273 in K18-hACE2 mice

We evaluated the protective activity of the mRNA-1273 vaccine against B.1.1.529 challenge. Although B.1.1.529 is less pathogenic in mice and hamsters (Bentley et al., 2021; Halfmann et al., 2022; Shuai et al., 2022), the virus still replicates to reasonably high levels (approximately 10-100 million copies of *N* gene/mg at 6 days post-infection (dpi)) in the lungs of K18-hACE2 mice (Halfmann et al., 2022). Five weeks after the second vaccine dose, mice were challenged via intranasal route with 10^4^ focus-forming units (FFU) of WA1/2020 D614G or B.1.1.529. Compared to the control mRNA vaccine, the 5 and 0.1 μg doses of mRNA-1273 vaccines prevented weight loss at 6 dpi after WA1/2020 D614G infection (**Fig 2A-B**). However, as B.1.1.529-challenged mice failed to lose weight, we could not use this metric to evaluate the protective activity of the mRNA-1273 vaccine.

**Figure 2.**
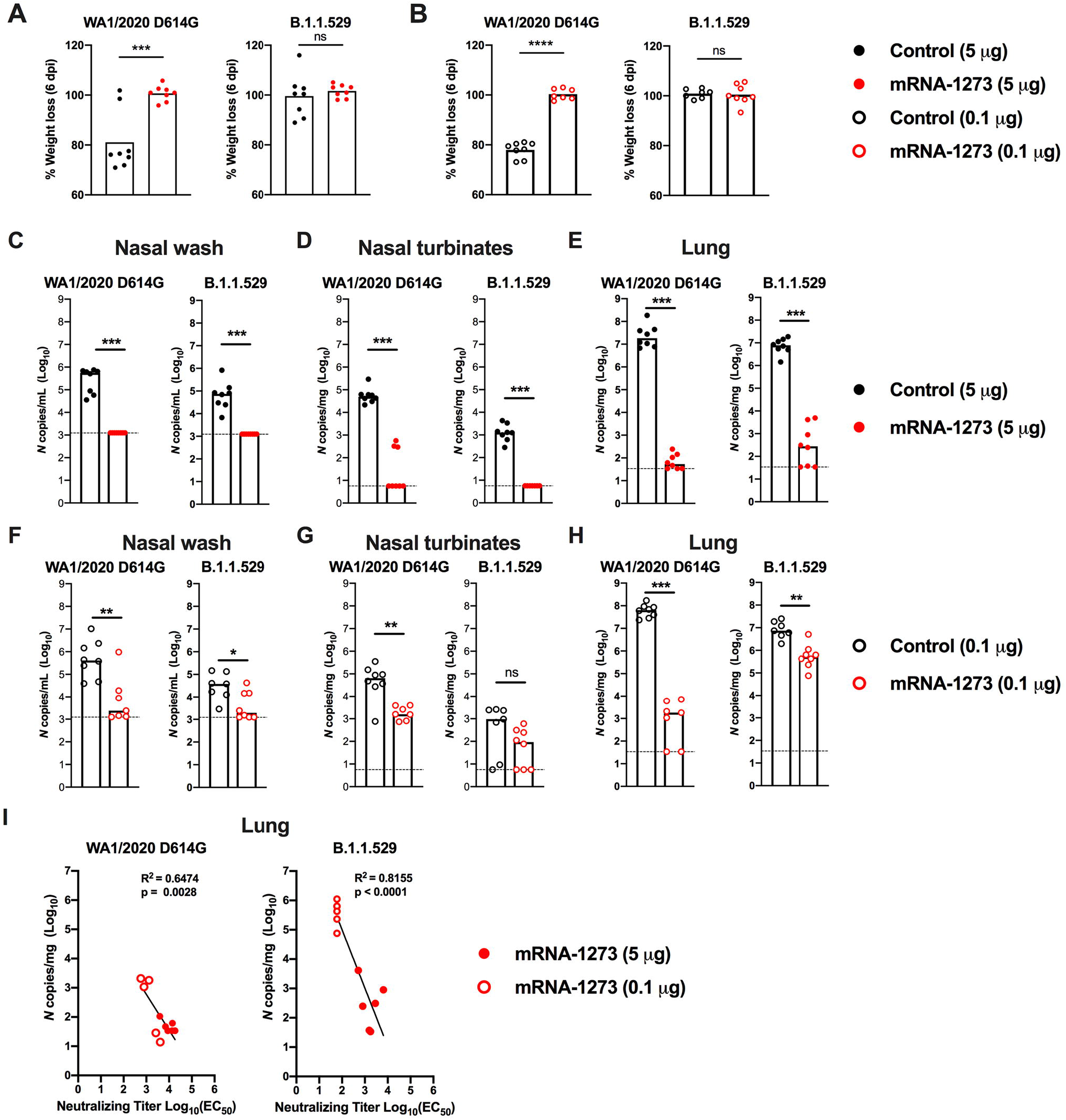
Protection against SARS-CoV-2 infection after mRNA vaccination in K18-hACE2 mice. Seven-week-old female K18-hACE2 transgenic mice were immunized with 5 or 0.1 μg of mRNA vaccines as described in **Fig 1A**. Five weeks after completing a primary vaccination series, mice were challenged with 10^4^ focus-forming units (FFU) of WA1/2020 D614G or B.1.1.529. **A-B**. Body weight change in animals immunized with 5 μg (**A**) or 0.1 μg (**B**) of control or mRNA-1273 vaccines between days 0 and 6 after challenge with WA1/2020 D614G or B.1.1.529. Data show mean values (n = 7-8, two experiments). **C-H**. Viral burden at 6 dpi in the nasal washes (**C, F**), nasal turbinates (**D, G**), and lungs (**E, H**) as assessed by qRT-PCR of the *N* gene after WA1/2020 D614G or B.1.1.529 challenge of mice immunized with 5 μg (**C-E**) or 0.1 μg (**F-H**) of control or mRNA-1273 vaccines (n = 7-8, two experiments, boxes illustrate median values, dotted lines show LOD). Statistical analyses: **A-B**, unpaired t test; **C-H**: Mann-Whitney test (ns, not significant; * *P* < 0.05; ** *P* < 0.01; *** *P* < 0.001; **** *P* < 0.0001). **I**. Correlation analyses comparing serum neutralizing antibody concentrations three weeks after the second vaccine dose and lung viral titers (6 dpi) in K18-hACE2 mice after challenge with WA1/2020 D614G (*left panel*) or B.1.1.529 (*right panel*); Pearson’s correlation R^2^ and *P* values are indicated as insets; closed symbols 5 μg vaccine dose; open symbols, 0.1 μg vaccine dose.

We next compared the levels of WA1/2020 D614G and B.1.1.529 infection in control mRNA-vaccinated K18-hACE2 mice at 6 dpi (**Fig 2C-H**). In the nasal washes of control mRNA-vaccinated K18-hACE2 mice, although some variability was observed, moderate amounts (10^5^ to 10^6^ copies of *N* per mL) of WA1/2020 D614G RNA were measured; approximately 10-fold lower levels (10^4^ to 10^5^ copies of *N* per mL) were measured after challenge with B.1.1.529 (**Fig 2C and F**). In the nasal turbinates, a similar pattern was seen with approximately 100-fold lower levels of B.1.1.529 RNA (~10^3^ versus 10^5^ copies of *N* per mg) (**Fig 2D and G**). In the lungs of control mRNA-vaccinated K18-hACE2 mice, approximately 10-fold less B.1.1.529 RNA was measured compared to WA1/2020 D614G RNA (**Fig 2E and H**).

We assessed the effects of mRNA-1273 vaccination on WA1/2020 D614G and B.1.1.529 infection in respiratory tract samples. The 5 μg dose of mRNA-1273 vaccine protected against WA1/2020 D614G infection with little viral RNA detected at 6 dpi (**Fig 2C-E**). In comparison, while B.1.1.529 viral RNA was not detected in the nasal washes or nasal turbinates of animals immunized with 5 μg of mRNA-1273, we observed breakthrough infection, albeit at low levels, in the lungs of most (5 of 8) animals (**Fig 2C-E**). Although the 0.1 μg dose of mRNA-1273 vaccine conferred protection against WA1/2020 D614G, several animals had viral RNA in nasal washes (4 of 7 mice), nasal turbinates (7 of 7), and lungs (5 of 7 mice). However, these breakthrough infections generally had reduced (100-100,000-fold) levels compared to the control mRNA vaccine (**Fig 2F-H**). After immunization with 0.1 μg of mRNA-1273, B.1.1.529 infection levels were lower in the nasal washes but not in the nasal turbinates (**Fig 2F-G**), in part due to the lower levels of viral RNA in the control mRNA-vaccinated samples. However, in the lungs, 7 of 7 mice vaccinated with the 0.1 μg dose sustained high levels of B.1.1.529 breakthrough infection with ~10^6^ copies of *N* per mg, although some protection still was observed (**Fig 2H**).

Serum neutralizing antibody titers showed an inverse correlation with amounts of viral RNA in the lung (**Fig 2I**) for both viruses, with more infection occurring in B.1.1.529-infected animals with lower neutralization titers. The correlation was most linear for B.1.1.529-challenged animals (R^2^ = 0.8155, *P* < 0.0001), with a minimum neutralizing titer of approximately 2,000 required to completely prevent infection at 6 dpi. Most of the breakthrough infections occurred with the lower 0.1 μg dose of mRNA vaccines, which models what might be occurring in immunocompromised or elderly individuals, or immunocompetent individuals at times remote from completion of their primary immunization series (Chen et al., 2021a; Choi et al., 2021; Evans et al., 2021).

We tested whether the mRNA-1273 vaccine could suppress cytokine and chemokine responses in the lung at 6 dpi of K18-hACE2 mice after challenge with WA1/2020 D614G or B.1.1.529 (**Fig 3A-B**). WA1/2020 D614G or B.1.1.529 infection of control mRNA vaccinated K18-hACE2 mice resulted in increased expression of several pro-inflammatory cytokines and chemokines including G-CSF, GM-CSF, IFNγ, IL-1β, IL-6, CXCL1, CXCL5, CXCL9, CXCL10, CCL2, CCL4, and TNF-α in lung homogenates (**Table S1 and S2**). Pro-inflammatory cytokine and chemokines in the lung at 6 dpi generally were lower in animals vaccinated with 5 μg dose of mRNA-1273 after challenge with WA1/2020 D614G or B.1.1.529 (**Fig 3A**). Although K18-hACE2 mice immunized with the lower 0.1 μg dose also showed diminished levels of cytokines and chemokines after WA1/2020 D614G infection, this protection was not observed after B.1.1.529 challenge; levels of pro-inflammatory cytokines in the lung were similar in mice immunized with the 0.1 μg dose control and mRNA-1273 vaccines after challenge with B.1.1.529 (**Fig 3B**).

**Figure 3.**
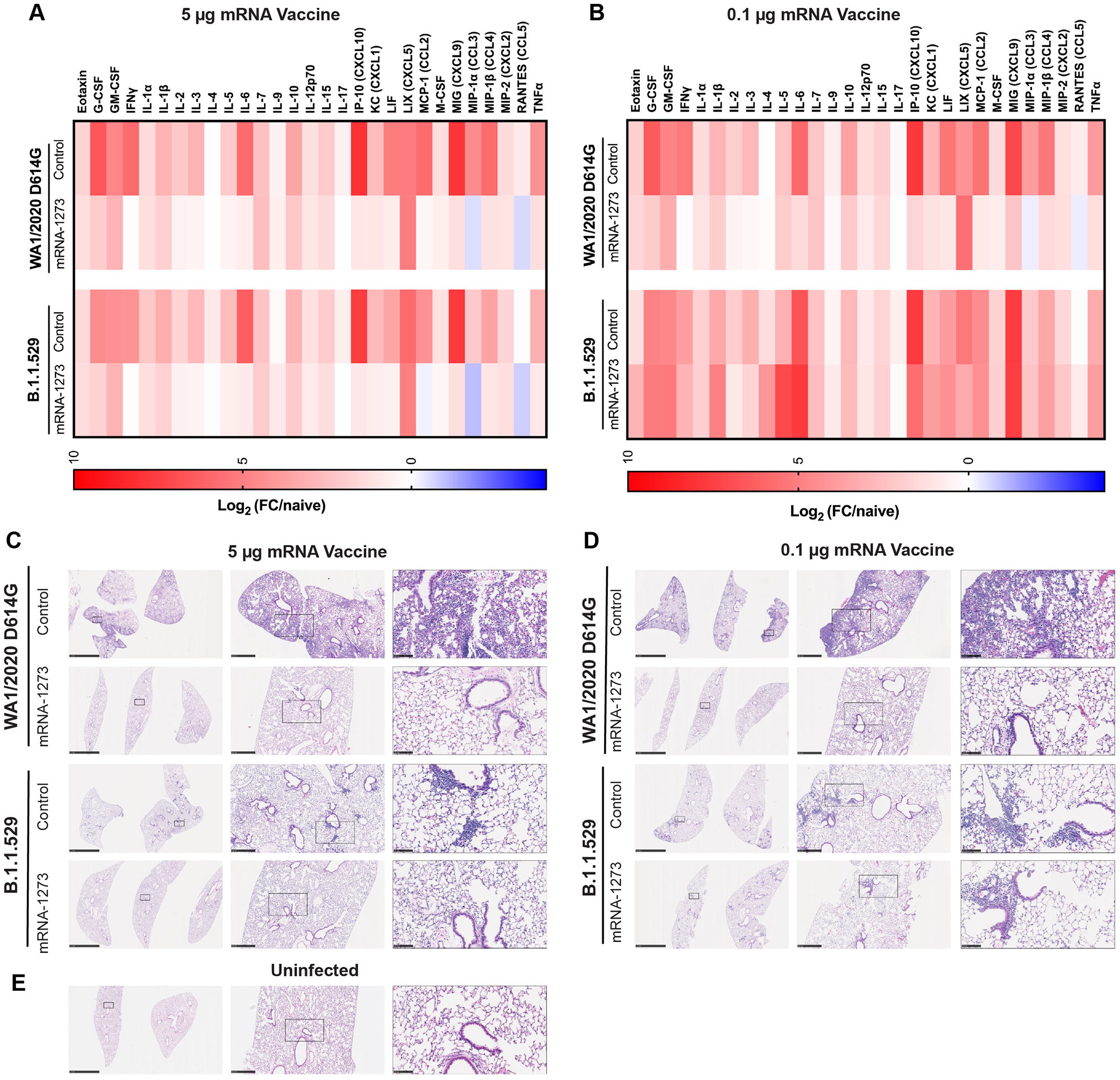
mRNA vaccine protection against disease in K18-hACE2 transgenic mice. Seven-week-old female K18-hACE2 transgenic mice were immunized with two 5 or 0.1 μg doses of mRNA vaccines, and challenged with WA1/2020 D614G or B.1.1.529. as described in **Fig 1 and 2**. **A-B**. Heat-maps of cytokine and chemokine levels in lung homogenates at 6 dpi in animals immunized with 5 μg (**A**) or 1 μg (**B**) doses of indicated mRNA vaccines. Fold-change was calculated relative to naive uninfected mice, and log_2_ values are plotted (2 experiments, n = 7-8 per group except naive, n = 4). The full data set is shown in **Table S1-S2**. **C-E**. Hematoxylin and eosin staining of lung sections harvested from control or mRNA-1273 vaccinated animals (5 μg dose, **C**; 1 μg dose, **D**) at 6 dpi with WA1/2020 D614G or B.1.1.529. A section from an uninfected animal (**E**) is shown for comparison. Images show low- (left; scale bars, 1 mm), moderate- (middle, scale bars, 200 μm), and high-power (bottom; scale bars, 50 μm). Representative images of multiple lung sections from n = 3 per group.

We performed histological analysis of lung tissues from immunized animals challenged with WA1/2020 D614G or B.1.1.529. Lung sections obtained at 6 dpi from mice immunized with either dose of control mRNA vaccine and challenged with WA1/2020 D614G showed severe pneumonia characterized by immune cell infiltration, alveolar space consolidation, vascular congestion, and interstitial edema (**Fig 3C-D**). In comparison, K18-hACE2 mice immunized with the control mRNA vaccines and challenged with B.1.1.529 showed less lung pathology, with focal airspace consolidation and immune cell infiltration, results that are consistent with the lower pathogenicity of B.1.1.529 in rodents (Abdelnabi et al., 2021; Bentley et al., 2021; Halfmann et al., 2022; Shuai et al., 2022).

Mice immunized with the high or low dose of mRNA-1273 and challenged with WA1/2020 D614G did not develop lung pathology, with histological findings similar to uninfected mice (**Fig 3C-E**). Mice immunized with the high dose of mRNA-1273 vaccine were protected against the mild pathological changes associated with B.1.1.529 infection (**Fig 3C**). However, animals immunized with the lower dose of mRNA-1273 showed similar lung pathology after B.1.1.529 infection as control mRNA-vaccinated animals, with patchy immune cell infiltration, airway space thickening, and mild alveolar congestion (**Fig 3D**).

### Effects of an mRNA-1273 booster dose on antibody responses and protection in K18-hACE2 mice

As booster doses are now used in humans to augment immunity and protection against variants, including B.1.1.529 (Atmar et al., 2022; Bar-On et al., 2021; Pajon et al., 2022), we evaluated their effects in K18-hACE2 mice. A cohort of 7-week-old female K18-hACE2 mice was immunized with a primary series over a 3-week interval with either a 5 or 0.25 μg dose of mRNA-1273 or a control vaccine (**Fig 4A**). The 0.25 μg dose used was higher than the 0.1 μg dose used above, in part due to the extended interval between the primary immunization series and boosting. After a 17 to 19-week rest period, blood was collected (pre-boost), animals were boosted with 1 μg of mRNA-1273 or a control vaccine, a second bleed (post-boost) was performed one month later (**Fig 4A**), and sera were tested for neutralizing activity. As expected, 5-to 10-fold lower pre-boost neutralizing titers were observed from animals immunized with the 0.25 μg than the 5 μg dose of mRNA-1273 (**Fig 4B-E and Fig S2**). While the titers against WA1/2020 D614G were high (GMT: 9,579, 5 μg; 3,096, 0.25 μg), we observed 10- to 20-fold lower levels (*P* < 0.01, 5 μg) of neutralizing antibodies against B.1.1.529, with half of the sera from mice vaccinated with the 0.25 μg formulation showing no inhibitory activity at the 1/60 limit of detection of the assay (**Fig 4B, D and Fig S2**). One month after boosting with mRNA-1273, serum neutralizing titers rose against both viruses (**Fig 4C, E and Fig S2**). All mice boosted with mRNA-1273 had neutralizing titers against B.1.1.529 above the estimated threshold (titer of 50) for protection (GMT: 6,124, 5 μg; 1,161, 0.25 μg).

**Figure 4.**
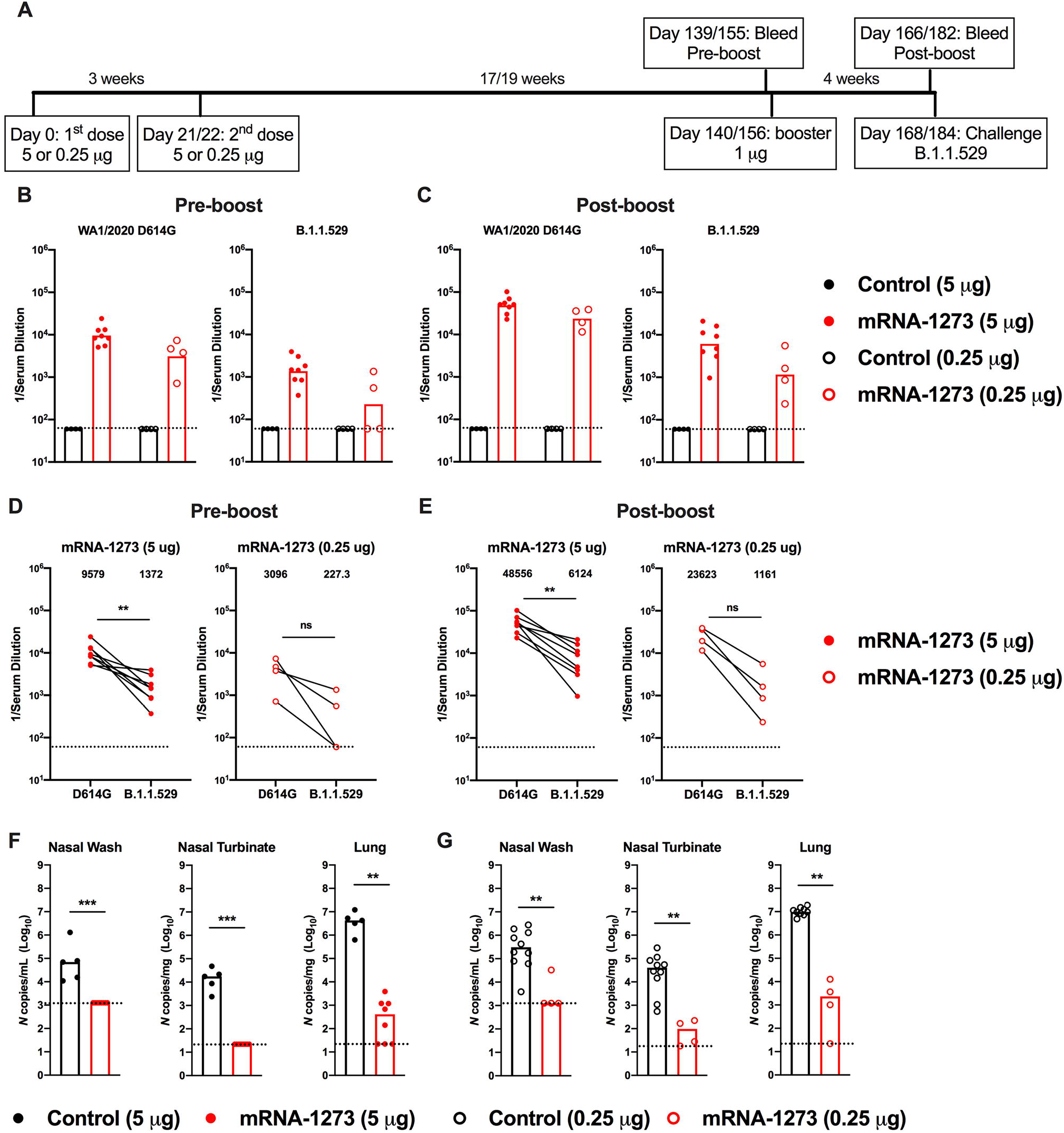
A booster dose of mRNA-1273 enhances neutralizing antibody responses and confers protection in K18-hACE2 mice. Seven-week-old female K18-hACE2 transgenic mice were immunized with 5 or 0.25 μg of mRNA vaccines and boosted 17 to 19 weeks later with 1 μg of mRNA-1273. **A**. Scheme of immunizations, blood draws, and virus challenge. **B-C**. Serum neutralizing antibody responses immediately before (**B**, pre-boost) and four weeks after (**C**, post-boost) a control or mRNA-1273 booster dose as judged by FRNT with WA1/2020 D614G (*left panel*) and B.1.1.529 (*right panel*) in mice immunized with 5 or 0.25 μg of control (n = 4) or mRNA-1273 (5 μg, n = 8; 0.25 μg, n =4) vaccines (one experiment, boxes illustrate geometric mean titer (GMT) values, dotted lines show the LOD). **D-E**. Paired analysis of pre-boost (**D**) and post-boost (**E**) serum neutralizing titers against WA1/2020 D614G and B.1.1.529 from individual mice (data from **B-C**) from samples obtained from animals that received a primary 5 μg (*left panel*) or 0.25 μg (*right panel*) dose series of mRNA-1273 vaccine (n = 4 to 8, one experiment, dotted lines show the LOD). GMT values are indicated at the top of the graphs. **F-G**. Four weeks after boosting with control or mRNA-1273, K18-hACE2 mice were challenged with 10^4^ FFU of B.1.1.529. Viral burden at 6 dpi in the nasal washes, nasal turbinates, and lungs as assessed by *N* gene levels in animals that had received a primary series immunization with 5 μg (**F**) or 0.25 μg (**G**) doses of control or mRNA-1273 vaccines (n = 7-8, two experiments, boxes illustrate median values, dotted lines show LOD). Statistical analyses. **D-E**. Wilcoxon signed-rank test; **F-G**: Mann-Whitney test (ns, not significant; ** *P* < 0.01; *** *P* < 0.001).

We evaluated the protective effects of mRNA-1273 boosting on B.1.1.529 infection by measuring viral RNA levels in nasal wash, nasal turbinates, and lungs at 6 dpi (**Fig 4F-G**). Compared to animals immunized and boosted with the control mRNA vaccine, K18-hACE2 mice vaccinated with the 5 or 0.25 μg primary series and boosted with mRNA-1273 showed reduced levels (*P* < 0.001) of B.1.1.529 viral RNA in the respiratory tract, with little to no detectable *N* gene copies in the nasal wash or turbinates. However, breakthrough infection, albeit at 2,500 to 6,000-fold lower levels (*P* < 0.001) than the control vaccine, was detected in the lungs of the majority (8 of 12) of mRNA-1273 boosted K18-hACE2 mice (**Fig 4F-G**). Thus, boosting of mice with mRNA-1273 vaccine, despite not being matched to the challenge virus, improves neutralizing antibody responses and reduces infection by B.1.1.529 in the upper and lower respiratory tracts.

### Immunogenicity of an Omicron-matched mRNA vaccine

As an alternative to a third dose of mRNA-1273, boosting with vaccines targeting the B.1.1.529 spike might provide enhanced immunity and protection. To begin to address this question, we generated a lipid-encapsulated mRNA vaccine (mRNA-1273.529) encoding a proline-stabilized SARS-CoV-2 spike from the B.1.1.529 virus. As a first test of its activity, we immunized BALB/c mice twice at 3-week intervals with 1 or 0.1 μg of mRNA 1273 or mRNA 1273.529 vaccines (**Fig 5A**). Three weeks after the first dose (Day 21) and two weeks after the second dose (Day 36), serum was collected (**Fig 5A**) and analyzed for binding to Wuhan-1 and B.1.1.529 spike proteins by ELISA (**Fig 5B**). At day 21 after the first dose, animals receiving 1 μg of mRNA-1273 showed approximately 11-fold higher levels of binding to homologous Wuhan-1 than heterologous B.1.1.529 spike, whereas equivalent responses to Wuhan-1 and B.1.1.529 spike proteins were detected after vaccination with 1 μg of mRNA-1273.529. In comparison, mice receiving the lower 0.1 μg dose of either mRNA-1273 or mRNA-1273.529 had low levels (near the limit of detection) of anti-spike antibody at day 21. Serum collected two weeks after the second dose of mRNA-1273 or mRNA-1273.529 also was tested for binding to Wuhan-1 and B.1.1.529 spike proteins. Immunization with either dose of mRNA-1273 resulted in higher (8 to 20-fold) serum IgG binding titers to Wuhan-1 than B.1.1.529 spike (**Fig 5B**). Reciprocally higher levels (3-fold) of serum IgG binding to B.1.1.529 than Wuhan-1 spike were seen after two 0.1 μg doses of mRNA-1273.529; however, immunization with two 1 μg doses of mRNA-1273.529 resulted in equivalent serum antibody binding to Wuhan-1 and B.1.1.529 spike proteins.

**Figure 5.**
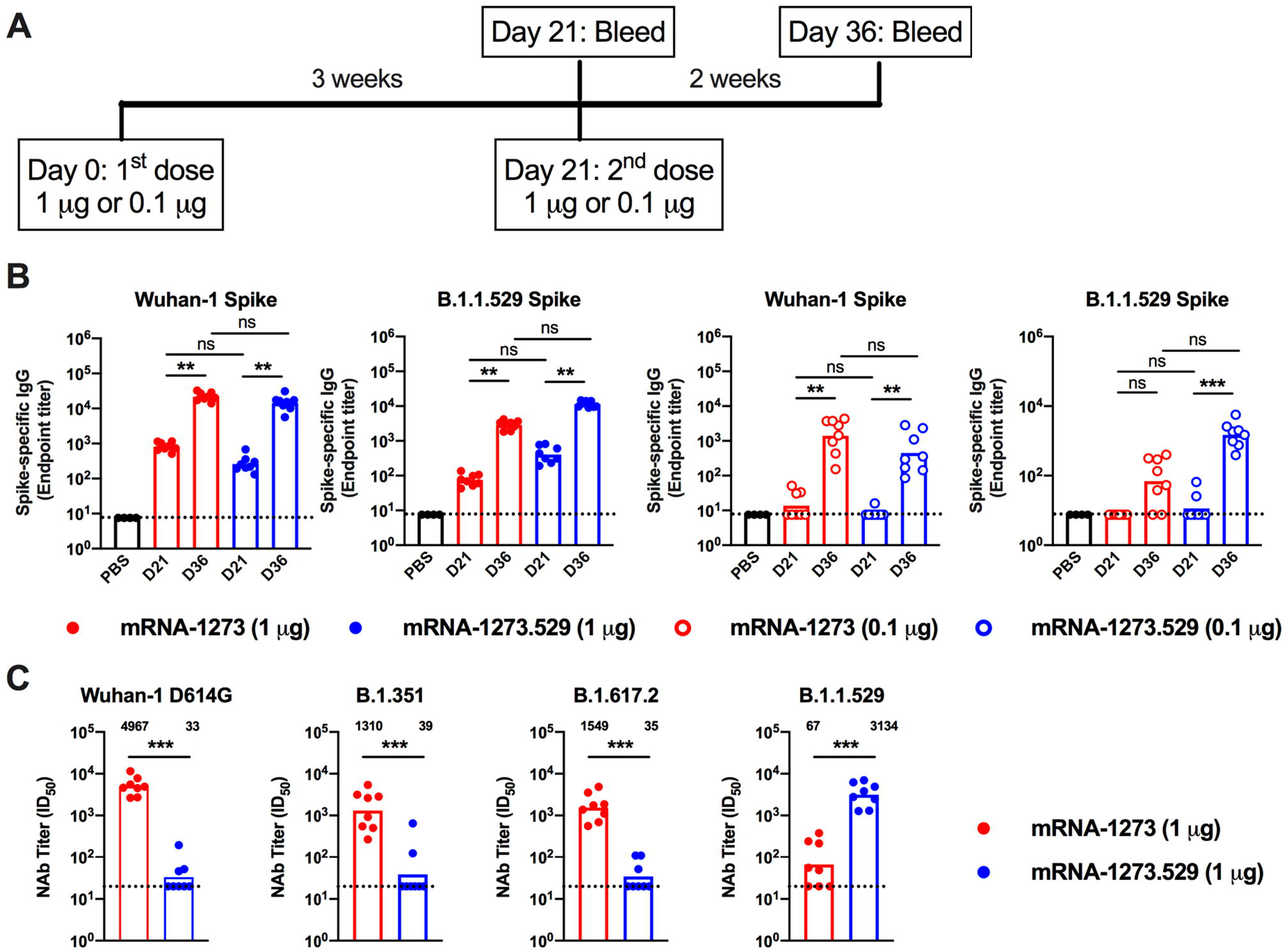
Antibody responses in BALB/c mice after immunization with mRNA-1273 and mRNA-1273.529 vaccines. Six-to-eight-week-old female BALB/c mice were immunized twice over a three-week interval with 1 μg of mRNA-1273 or mRNA-1273.529 vaccine or a PBS control (black circles). Immediately before (Day 21) or two weeks after (Day 36) the second vaccine dose, serum was collected. **A**. Scheme of immunization and blood draws. **B**. Serum antibody binding to Wuhan-1 or B.1.1.529 spike proteins by ELISA (n = 8, two experiments, boxes illustrate mean values, dotted lines show the LOD). **C**. Neutralizing activity of serum obtained two weeks after (Day 36) immunization with mRNA-1273 or mRNA-1273.529 vaccine against VSV pseudoviruses displaying the spike proteins of Wuhan-1 D614G, B.1.351 (Beta), B.1.617.2 (Delta), or B.1.1.529 (Omicron) (n = 8, two experiments, boxes illustrate geometric mean values, dotted lines show the LOD). GMT values are indicated above the columns.

We next tested the inhibitory activity of serum antibodies of BALB/c mice that received two 1 μg doses of mRNA-1273 or mRNA-1273.529 using a vesicular stomatitis virus (VSV)-based pseudovirus neutralization assay and spike proteins of Wuhan-1 D614G, B. 1.351, B.1.617.2, or B.1.1.529 (**Fig 5C**). Two weeks after the second vaccine dose, animals immunized with mRNA-1273 had high serum neutralizing titers against Wuhan-1 D614G (GMT: 4,967) with approximately four-fold reductions (*P* < 0.01) against viruses displaying B.1.351 (GMT: 1,310) or B.1.617.2 (GMT: 1,549). However, and consistent with data in K18-hACE2 mice (**Fig 4D**) and humans (Dejnirattisai et al., 2022; Liu et al., 2021a), BALB/c mice vaccinated with mRNA-1273 had substantially lower (74-fold reduced, GMT:67, *P* < 0.01) neutralizing titers against B.1.1.529 with several samples falling near the presumed 1/50 threshold of protection. In comparison, animals immunized with two 1 μg doses of the Omicron variant-targeted mRNA-1273.529 vaccine showed a distinct pattern. Although high neutralization titers were observed against B.1.529 (GMT: 3,314), lower levels of neutralization (85 to 100-fold less, *P* < 0.01) were detected against Wuhan-1 D614G (GMT: 33), B.1.351 (GMT: 39), and B.1.617.2 (GMT: 35), with most of the samples at the limit of detection of the neutralizing assay. Thus, and as suggested recently in another preliminary study (Lee et al., 2022), a primary vaccination series with an Omicron-specific mRNA vaccine induces potently neutralizing antibodies against Omicron but not against historical viruses or other SARS-CoV-2 variants.

### Protective effects of an Omicron-targeted mRNA vaccine

As BALB/c and K18-hACE2 mice immunized with a primary series were unavailable, we took advantage of an existing cohort of female 129S2 mice that had been immunized with a primary series over a 3-week interval with either a 5 or 0.25 μg dose of mRNA-1273 or a control mRNA vaccine and then rested for 10 to 11 weeks (**Fig 6A**). Blood was collected (pre-boost sample), and groups of animals were boosted homologously or heterologously with 1 μg of control mRNA, mRNA-1273, or mRNA-1273.529 vaccine, the same dose used in our BALB/c mouse studies above. Three to four weeks later, a second post-boost blood sample was collected (**Fig 6A**), and the neutralizing activity of pre- and post-boost serum antibodies were measured and compared. Mice that received 5 or 0.25 μg doses of mRNA-1273 had high-levels (GMT: 29,161, 5 μg; 5,749, 0.25 μg) of pre-boost neutralizing antibodies against WA1/2020 N501Y/D614G (**Fig 6B and S3**). We used the WA1/2020 N501Y/D614G strain because the N501Y substitution enables engagement of murine ACE2 and productive infection of conventional strains of laboratory mice (Gu et al., 2020; Liu et al., 2021b; Rathnasinghe et al., 2021; Ying et al., 2021). However, mice that received 5 or 0.25 μg doses of mRNA-1273 had 15- to 18-fold lower serum pre-boost neutralizing titers against B.1.1.529 (GMT: 1891, 5 μg; 317, 0.25 μg; *P* < 0.0001) (**Fig 6B and S3**). One month after boosting with mRNA-1273 or mRNA-1273.529, neutralizing titers against WA1/2020 N501Y/D614G were approximately 2 to 6-fold-higher than before boosting (**Fig 6C-D and S3**). In comparison, boosting with mRNA-1273.529 resulted in 22 to 42-fold (*P* < 0.01) higher neutralizing titers against B.1.1.529, whereas the mRNA-1273 booster increased titers against B.1.1.529 by 2 to 5-fold (*P* < 0.05) (**Fig 6C, E and Fig S3**). Thus, while both mRNA-1273 and mRNA-1273.529 boosters augmented serum neutralizing activity of B.1.1.529, an Omicron-matched vaccine produced higher titers of neutralizing antibodies.

**Figure 6.**
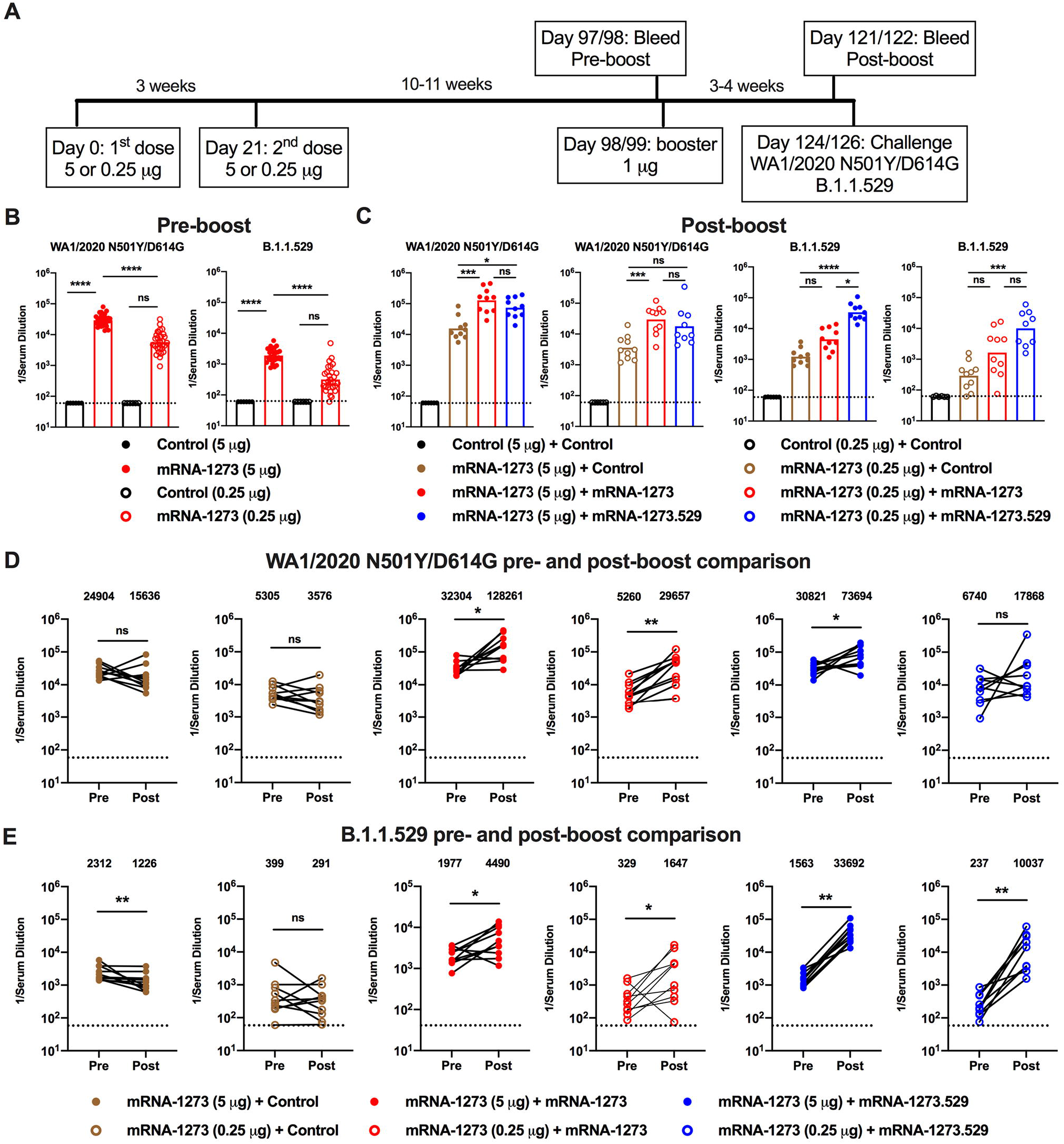
Booster doses of mRNA-1273 or mRNA-1273.529 enhance neutralizing antibody responses in 129S2 mice. Seven-week-old female 129S2 mice were immunized with 5 or 0.25 μg of mRNA vaccines and then boosted 10 to 11 weeks later with 1 μg of control mRNA, mRNA-1273, or mRNA-1273.529. **A**. Scheme of immunizations, blood draws, and virus challenge. **B-C**. Serum neutralizing antibody responses immediately before (**B**, pre-boost) and three to four weeks after (**C**, post-boost) a control, mRNA-1273, or mRNA-1273.529 booster dose as judged by FRNT with WA1/2020 N501Y/D614G (*left panel(s)*) and B.1.1.529 (*right panel(s)*) in 129S2 mice that received primary series immunizations with 5 or 0.25 μg of control (n = 6) or mRNA-1273 (n = 30) vaccines (two experiments, boxes illustrate geometric mean values, dotted lines show the LOD). **D-E**. Paired analysis of pre- and post-boost serum neutralizing titers against WA1/2020 D614G (**D**) and B.1.1.529 (**E**) viruses from samples obtained from animals (data from **B-C**) that received the following primary and booster immunizations: mRNA-1273 (5 or 0.25 μg) + control booster, mRNA-1273 (5 or 0.25 μg) + mRNA-1273 booster, mRNA-1273 (5 or 0.25 μg) + mRNA-1273.529 booster (n = 10, two experiments, dotted lines show the LOD). GMT values are indicated at the top of the graphs. Statistical analyses. **B-C**: One-way ANOVA with Dunn’s post-test; **D-E**. Wilcoxon signed-rank test (ns, not significant; * *P* < 0.05; ** *P* < 0.01; *** *P* < 0.001; **** *P* < 0.0001).

Three or four days after the post-boost bleed, 129S2 mice were challenged by the intranasal route with 10^5^ FFU of WA1/2020 N501Y/D614G or B.1.1.529, and viral RNA levels at 3 dpi were measured in the nasal washes, nasal turbinates, and lungs. B.1.1.529 is less pathogenic in 129S2 mice (Halfmann et al., 2022) with ~100-fold lower levels of viral RNA in the upper and lower respiratory tract than after WA1/2020 N501YD614G infection (**Fig 7A-B**). Nonetheless, substantial viral replication occurred allowing for evaluation of vaccine protection. Mice vaccinated with either 5 or 0.25 μg doses of mRNA-1273 and boosted with either mRNA-1273 or mRNA-1273.529 showed almost complete protection against WA1/2020 N501Y/D614G infection in the nasal washes, nasal turbinates and lungs, with approximately 100,000- to 1,000,000-fold reductions in viral RNA levels compared to mice immunized with control mRNA vaccine. Mice primed with the 5 μg doses of mRNA-1273 and boosted with either mRNA-1273 or mRNA-1273.529 analogously showed robust and equivalent protection against B.1.1.529 infection (**Fig 7A**). In comparison, animals primed with the lower 0.25 μg dose of mRNA-1273 showed some differences after boosting and B.1.1.529 challenge (**Fig 7B**). Although B.1.1.529 viral RNA levels were reduced in upper respiratory tract tissues after boosting with either mRNA-1273 or mRNA-1273.529, there was a trend (*P* = 0.07) toward lower levels in the nasal turbinates in animals boosted with mRNA-1273.529. Moreover, a 27-fold reduction (*P* < 0.01) in B.1.1.529 infection in the lungs was observed in mice boosted with the Omicron-matched mRNA-1273.529 compared to the mRNA-1273 vaccine.

**Figure 7.**
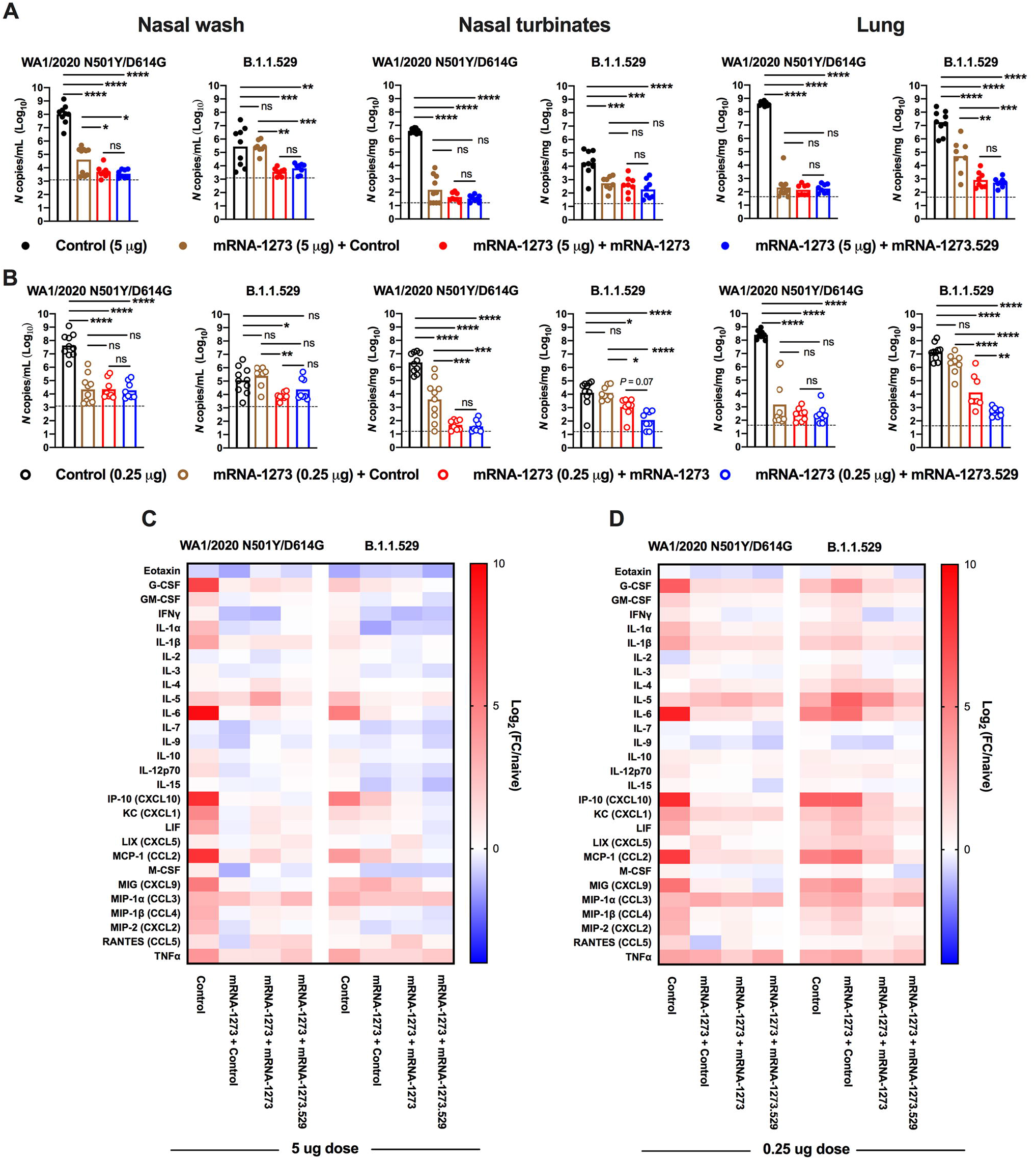
Booster doses of mRNA-1273 or mRNA-1273.529 enhance protection against B.1.1.529 infection in 129S2 mice. Seven-week-old female 129S2 mice were immunized with 5 or 0.25 μg of mRNA vaccines, boosted with 1 μg of control mRNA, mRNA-1273, or mRNA-1273.529 and challenged with WA1/2020 N501Y/D614G or B.1.1.529, as described in **Fig 6. A-B**. Viral RNA levels at 3 dpi in the nasal washes, nasal turbinates, and lungs after WA1/2020 N501Y/D614G or B.1.1.529 challenge of mice immunized with 5 μg (**A**) or 0.25 μg (**B**) of control or mRNA-1273 vaccines and boosted with control, mRNA-1273, or mRNA-1273.529 vaccine (n = 8-10 per group, two experiments, boxes illustrate mean values, dotted lines show LOD; One-way ANOVA with Tukey’s post-test: ns, not significant; * *P* < 0.05; ** *P* < 0.01; *** *P* < 0.001; **** *P* < 0.0001). **C-D**. Heat-maps of cytokine and chemokine levels in lung homogenates at 3 dpi with WA1/2020 N501Y/D614G or B.1.1.529 in animals immunized with 5 μg (**C**) or 0.25 μg (**D**) doses of control or mRNA-1273 vaccines and then boosted with control, mRNA-1273, or mRNA-1273.529 vaccines. Fold-change was calculated relative to naive mice, and log_2_ values are plotted (2 experiments, n = 8 per group except naive, n = 4). The full data set is shown in **Table S3-S4**.

As an independent metric of protection, we measured cytokine and chemokine levels in lung homogenates of the vaccinated and challenged 129S2 mice at 3 dpi (**Fig 7C-D and Table S3 and S4**). Mice immunized with 5 or 0.25 μg doses of mRNA-1273 and then given a booster dose of either control, mRNA-1273, or mRNA-1273.529 vaccines generally showed lower levels of pro-inflammatory cytokines and chemokines after WA1/2020 N501Y/D614G infection than animals that received three doses of control mRNA vaccine. In comparison, after B.1.1.529 challenge several differences were noted: (a) the inflammatory response after B.1.1.529 infection was lower in magnitude in control mRNA-vaccinated mice than after WA1/2020 N501Y/D614G infection, consistent with its less pathogenic nature in 129 mice (Halfmann et al., 2022); (b) animals immunized with 5 μg doses of mRNA-1273 and then boosted with control, mRNA-1273, or mRNA-1273.529 vaccines showed reduced levels of cytokines and chemokines compared to those receiving three doses of control mRNA, indicating protection against inflammation by homologous or heterologous boosting (**Fig 7C**); and (c) mice immunized with two 0.25 μg doses of mRNA-1273 and boosted with control vaccine showed little reduction in cytokine and chemokine levels compared to mice receiving three doses of control mRNA vaccine following B.1.1.529 challenge (**Fig 7D**). While boosting with mRNA-1273 reduced the levels of most lung inflammatory mediators, the effect was greater in animals boosted with mRNA-1273.529. Thus, and consistent with the virological data, protection against B.1.1.529-induced lung inflammation was modestly enhanced in animals boosted with the Omicron-matched mRNA-1273.529 vaccine.

## DISCUSSION

Vaccine-induced immunity against SARS-CoV-2 can limit human disease and curtail the COVID-19 global pandemic. The emergence of SARS-CoV-2 variants with constellations of amino acid changes in the NTD and RBD of the spike protein jeopardizes vaccines designed against historical SARS-CoV-2 strains. In the current study, we evaluated in mice the protective activity of high and low dose formulations of Moderna mRNA-1273 vaccine and matched (mRNA-1273.529) and non-matched (mRNA-1273) booster doses against the B.1.1.529 Omicron variant. Although the low-dose vaccine arms were designed to model individuals with suboptimal immune responses, they induce levels of neutralizing antibody in mice comparable to those measured in human sera after completion of a primary two-dose vaccination series with mRNA-1273 (Anderson et al., 2020; Widge et al., 2021; Wu et al., 2021). Immunization of mice with the high-dose formulation of mRNA-1273 induced neutralizing antibodies that inhibited infection in cell culture of both the historical WA1/2020 D614G and B.1.1.529 viruses, although we observed reduced efficacy against the Omicron variant, as seen with human sera (Cameroni et al., 2021; Cao et al., 2021; Dejnirattisai et al., 2022; Liu et al., 2021a). Challenge studies in K18-hACE2 and 129S2 mice showed robust protection against both SARS-CoV-2 strains with the high-dose primary series of mRNA-1273 vaccine. In comparison, the neutralizing antibodies induced by the low-dose series of mRNA-1273 vaccine showed less inhibitory activity against B.1.1.529, which correlated with breakthrough infection in the upper and lower respiratory tracts. Analysis of cytokines and histology corroborated the low to minimal protection against B.1.1.529 in K18-hACE2 mice by the low-dose series of mRNA-1273 vaccine.

We extended these studies by evaluating the effects of matched and non-matched mRNA vaccine boosters on antibody responses and protection. In a study with a small cohort of K18-hACE2 mice, we observed increased serum neutralizing titers one month after boosting with mRNA-1273, although the response to B.1.1.529 was lower than against WA1/2020. These data are consistent with data from humans who received a primary mRNA-1273 series and homologous booster; while neutralizing titers were increased against B.1.1.529 after boosting, levels were lower than against WA1/2020 D614G and waned faster (Pajon et al., 2022). Because of this pattern, some have advocated for the development of an Omicron variant-targeted booster to improve protection against infection, disease, and transmission (Waltz, 2022). In our studies in BALB/c mice, as a primary immunization series, an Omicron-targeted mRNA-1273-529 vaccine robustly induced neutralizing antibodies in serum against B.1.1.529. However, neutralizing antibody titers against heterologous WA1/2020 D614G, B. 1.351, and B.1.617.2 strains were much lower, as reported by another group (Lee et al., 2022) and also described in unvaccinated humans that experienced a primary B.1.1.529 infection (Rössler et al., 2022). Thus, in unvaccinated, uninfected individuals, bivalent or multivalent mRNA vaccines (Pajon et al., 2022; Ying et al., 2021) (*e.g*., mRNA-1273 + mRNA-1273.529) might be required to achieve necessary breadth of humoral responses.

When mRNA-1273.529 was administered as a booster dose after a mRNA-1273 primary series, we observed enhanced neutralizing responses against B.1.1.529, and this was associated with protection against B.1.1.529 infection in the upper and lower respiratory tracts. Moreover, boosting with mRNA-1273.529 also enhanced neutralizing antibody responses against the historical WA1/2020 D614G strain, albeit less so, which suggests no substantive detriment in breadth of the mRNA-1273.529 response in the setting of boosting in contrast to the primary immunization series. Notwithstanding these results, when compared to boosting with the historical mRNA-1273 vaccine, which increased neutralizing antibody responses against B.1.1.529 to a lesser degree, the differences in virological protection in the upper and lower airway were noticeably modest. When mice were given a high-dose mRNA-1273 primary vaccination series, there was no virological benefit of the Omicron-matched booster compared to the mRNA-1273 booster, which agrees with recent data from non-human primates (Gagne et al., 2022). However, when mice given a low-dose primary vaccination series, protection against lung infection and inflammation was slightly greater in animals boosted with the Omicron-matched mRNA-1273.529 vaccine. The relative disparity between the larger differences in boosting of neutralizing antibodies by mRNA-1273 and mRNA-1273.529 vaccines and the smaller effects on infection at one month post-boost could reflect the protective effects of pre-existing or anamnestic cross-reactive T cells (Gao et al., 2022; Keeton et al., 2022) or Fc effector function and immune cell engagement by cross-reactive non-neutralizing antibodies (Bartsch et al., 2021). Further studies examining the durability of protection afforded by mRNA-1273 versus Omicron-matched mRNA-1273.529 boosters are warranted.

### Limitations of study

We note several limitations in our study. (1) Female K18-hACE2 and 129S2 mice were used to allow for group caging, and some of our studies had smaller cohorts and were performed in different strains, due to animal availability. Follow-up experiments in male mice and with larger cohorts are needed to confirm and extend these results. (2) We used a B.1.1.529 Omicron isolate that lacks an R346K mutation; this substitution or ones in the emerging BA.2 lineage might further affect vaccine-induced virus neutralization and protection. (3) Our analysis focused on antibody responses and did not account for possible cross-reactive T cell responses, which could impact protective immunity. (4) The B.1.1.529 virus is less pathogenic in mice (Abdelnabi et al., 2021; Bentley et al., 2021; Halfmann et al., 2022). This could lead to an overestimation of protection compared to other more virulent strains in mice. (5) We analyzed mRNA-1273 and mRNA-1273.529 booster responses and protection one month after administration. A time course analysis is needed to assess the durability of the enhanced neutralizing antibody responses and protection against B.1.1.529. (6) Experiments were performed exclusively in mice to allow for analysis of multiple arms and comparisons. Vaccination, boosting, and B.1.1.529 challenge studies in other animal models and ultimately humans will be required for corroboration.

In summary, our studies in mice show protection against B.1.1.529 infection and disease when mRNA-1273 or Omicron-matched mRNA-1273.529 boosters are administered. While the low-dose primary immunization series of mRNA-1273 protected against WA1/2020 challenge, we observed substantial loss of serum neutralizing activity against B.1.1.529, and this was associated with breakthrough infection, inflammation, and disease in the lung after B.1.1.529 challenge. Despite diminished cross-variant neutralization responses by Omicron-matched and historical mRNA vaccines when administered as primary series immunizations, their delivery as boosters improved neutralizing responses against B.1.1.529, with slightly better protection conferred by the matched mRNA-1273.529 booster. Boosting with historical vaccines, variant-matched mRNA vaccines (Choi et al., 2021; Ying et al., 2021) or possibly heterologous platforms targeting historical spike proteins (*e.g.*, adenoviral vectored or protein subunit vaccines (Atmar et al., 2022)) could minimize B.1.1.529 breakthrough infections by increasing the magnitude of neutralizing anti-SARS-CoV-2 antibodies (Atmar et al., 2021; Munro et al., 2021) or expanding the breadth of the antibody repertoire to better control variant strains (Falsey et al., 2021; Naranbhai et al., 2022; Wang et al., 2021). While our data suggest that certain cohorts (those with lower starting antibody responses) might benefit more from an Omicron-matched vaccine, further studies evaluating the magnitude and durability of the boosted immune responses are needed, especially in key vulnerable populations including the elderly, immunocompromised and immunosuppressed.

## Acknowledgements

This study was supported by the NIH (R01 AI157155, NIAID Centers of Excellence for Influenza Research and Response (CEIRR) contracts HHSN272201400008C, 75N93021C00014, and 75N93019C00051). We thank Marciela DeGrace for help in study design and funding support, Pei-Yong Shi for the WA1/2020 strains, and Peter Halfmann and Yoshihiro Kawaoka for the B.1.1.529 isolate used in this study. We also acknowledge the Pulmonary Morphology Core at Washington University School of Medicine for tissue sectioning and slide imaging, and thank Michael Whitt for support on VSV-based pseudovirus production.

## Author contributions

B.Y. and S.M.S. performed and analyzed live virus neutralization assays. B.Y., S.M.S., B.W., C.Y.L., O.K., S.M., Z.C. and J.B.T. performed mouse experiments. B.W. and B.Y. performed and analyzed viral burden analyses. B.Y., L.M., T.K., C.S., and A.W. performed ELISA binding experiments and analysis. K.W., D.L., D.M.B., and L.E.A. performed pseudovirus neutralization assays and analysis. A.C., S.E. and D.K.E. provided mRNA vaccines and helped design experiments. L.B.T. and M.S.D. designed studies and supervised the research. M.S.D. and L.B.T. wrote the initial draft, with the other authors providing editorial comments.

## SUPPLEMENTAL FIGURE LEGENDS

**Figure S1. Serum neutralization of WA1/2020 D614G and B.1.1.529, Related to Fig 1.** Seven-week-old female K18-hACE2 transgenic mice were immunized with 5 or 0.1 μg of mRNA vaccines. Serum neutralizing antibody responses against WA1/2020 D614G and B.1.1.529 were assessed three weeks after the second vaccine dose (control mRNA, *top*; mRNA-1273, *bottom*) from mice immunized with 5 or 0.1 μg of control (n = 4) or mRNA-1273 (n = 12) vaccines. Neutralization curves corresponding to individual mice are shown for the indicated vaccines. Each point represents the mean of two technical replicates.

**Figure S2. Serum neutralization of WA1/2020 D614G and B.1.1.529, Related to Fig 4.** Seven-week-old female K18-hACE2 transgenic mice were immunized with 5 or 0.25 μg of mRNA vaccines and then boosted approximately 17 to 19 weeks later with 1 μg of mRNA-1273. Serum neutralizing antibody responses against WA1/2020 D614G and B.1.1.529 immediately before (**A**, pre-boost) and four weeks after (**B**, post-boost) a control or mRNA-1273 booster dose from mice immunized with 5 or 0.25 μg of control (n = 4) or mRNA-1273 (5 μg, n = 8; 0.25 μg, n =4) vaccines. Neutralization curves corresponding to individual mice are shown for the indicated immunizations. Each point represents the mean of two technical replicates.

**Figure S3. Pre- and post-boost serum neutralization of WA1/2020 N501Y/D614G and B.1.1.529, Related to Fig 6.** Seven-week-old female 129S2 mice were immunized with 5 or 0.25 μg of mRNA vaccines and then boosted 10 to 11 weeks later with 1 μg of control mRNA, mRNA-1273, or mRNA-1273.529. Neutralizing antibody responses against WA1/2020 N501Y/D614G and B.1.1.529 from serum immediately before (top) or one month after boosting (bottom) with indicated vaccines from 129S2 mice that had received primary series immunizations with 5 or 0.25 μg of control or mRNA-1273 vaccines. Neutralization curves corresponding to individual mice are shown for the indicated immunizations. Serum are from two independent experiments, and each point from a neutralization curve represents the mean of two technical replicates.

## SUPPLEMENTAL TABLE TITLES

**Supplemental Table S1. Cytokine and chemokine concentrations in K18-hACE2 mice vaccinated with two 5 μg doses of mRNA vaccines and challenged with WA1/2020 D614G or B.1.1.529, Related to Fig 3.**

**Supplemental Table S2: Cytokine and chemokine concentrations in K18-hACE2 mice vaccinated with two 0.1 μg doses of mRNA vaccines and challenged with WA1/2020 D614G or B.1.1.529, Related to Fig 3.**

**Supplemental Table S3: Cytokine and chemokine concentrations in 129S2 mice immunized with 5 μg primary series and 1 μg booster dose and challenged with WA1/2020 N501Y/D614G or B.1.1.529, Related to Fig 7.**

**Supplemental Table S4: Cytokine and chemokine concentrations in 129S2 mice immunized with 0.25 μg primary series and 1 μg booster dose and challenged with WA1/2020 N501Y/D614G or B.1.1.529, Related to Fig 7.**

## STAR METHODS

### RESOURCE AVAILABILITY

#### Lead contact

Further information and requests for resources and reagents should be directed to the Lead Contact, Michael S. Diamond.

#### Materials availability

All requests for resources and reagents should be directed to the Lead Contact author. This includes viruses, vaccines, and primer-probe sets. All reagents will be made available on request after completion of a Materials Transfer Agreement (MTA). The mRNA vaccines (control, mRNA-1273, and mRNA-1273.529) can be obtained under an MTA with Moderna (contact: Darin Edwards).

#### Data and code availability

All data supporting the findings of this study are available within the paper and are available from the corresponding author upon request. This paper does not include original code. Any additional information required to reanalyze the data reported in this paper is available from the lead contact upon request.

### EXPERIMENTAL MODEL AND SUBJECT DETAILS

#### Cells

African green monkey Vero-TMPRSS2 (Zang et al., 2020) and Vero-hACE2-TMPRRS2 (Chen et al., 2021c) cells were cultured at 37°C in Dulbecco’s Modified Eagle medium (DMEM) supplemented with 10% fetal bovine serum (FBS), 10□mM HEPES pH 7.3, 1□mM sodium pyruvate, 1× non-essential amino acids, and 100□U/mL of penicillin-streptomycin. Vero-TMPRSS2 cells were supplemented with 5 μg/mL of blasticidin. Vero-hACE2-TMPRSS2 cells were supplemented with 10 μg/mL of puromycin. All cells routinely tested negative for mycoplasma using a PCR-based assay.

#### Viruses

The WA1/2020 recombinant strain with D614G substitution was described previously (Plante et al., 2020). The B.1.1.529 isolate (hCoV-19/USA/WI-WSLH-221686/2021) was obtained from an individual in Wisconsin as a midturbinate nasal swab and passaged once on Vero-TMPRSS2 cells (Imai et al., 2020). All viruses were subjected to next-generation sequencing (Chen et al., 2021c) to confirm the introduction and stability of substitutions. All virus experiments were performed in an approved biosafety level 3 (BSL-3) facility.

#### Mice

Animal studies were carried out in accordance with the recommendations in the Guide for the Care and Use of Laboratory Animals of the National Institutes of Health. For studies (K18-hACE2 and 129S2 mice) at Washington University School of Medicine, the protocols were approved by the Institutional Animal Care and Use Committee at the Washington University School of Medicine (assurance number A3381–01). Virus inoculations were performed under anesthesia that was induced and maintained with ketamine hydrochloride and xylazine, and all efforts were made to minimize animal suffering. For studies with BALB/c mice, animal experiments were carried out in compliance with approval from the Animal Care and Use Committee of Moderna, Inc. Sample size for animal experiments was determined on the basis of criteria set by the institutional Animal Care and Use Committee. Experiments were neither randomized nor blinded.

Heterozygous K18-hACE2 C57BL/6J mice (strain: 2B6.Cg-Tg(K18-ACE2)2Prlmn/J, Cat # 34860) were obtained from The Jackson Laboratory. 129S2 mice (strain: 129S2/SvPasCrl, Cat # 287) and BALB/c mice (strain: BALB/cAnNCrl, Cat # 028) were obtained from Charles River Laboratories. Animals were housed in groups and fed standard chow diets.

### METHOD DETAILS

#### Pre-clinical vaccine mRNA and lipid nanoparticle production process

A sequence-optimized mRNA encoding prefusion-stabilized Wuhan-Hu-1 (mRNA-1273) SARS-CoV-2 S-2P or B.1.1.529 (mRNA-1273.529) protein was synthesized *in vitro* using an optimized T7 RNA polymerase-mediated transcription reaction with complete replacement of uridine by N1m-pseudouridine (Nelson et al., 2020). The pre-clinical mRNA-1273.529 vaccine encoded the following substitutions: A67V, Δ69-70, T95I, G142D, Δ143-145, Δ211, L212I, ins214EPE, G339D, S371L, S373P, S375F, K417N, N440K, G446S, S477N, T478K, E484A, Q493K, G496S, Q498R, N501Y, Y505H, T547K, D614G, H655Y, N679K, P681H, N764K, D796Y, N856K, Q954H, N969K, and L981F. A non-translating control mRNA was synthesized and formulated into lipid nanoparticles as previously described (Corbett et al., 2020). The reaction included a DNA template containing the immunogen open-reading frame flanked by 5’ untranslated region (UTR) and 3’ UTR sequences, and was terminated by an encoded polyA tail. After RNA transcription, the cap-1 structure was added using the vaccinia virus capping enzyme and 2⍰-*O*-methyltransferase (New England Biolabs). The mRNA was purified by oligo-dT affinity purification, buffer exchanged by tangential flow filtration into sodium acetate, pH 5.0, sterile filtered, and kept frozen at –20°C until further use.

The mRNA was encapsulated in a lipid nanoparticle through a modified ethanol-drop nanoprecipitation process described previously (Hassett et al., 2019). Ionizable, structural, helper, and polyethylene glycol lipids were briefly mixed with mRNA in an acetate buffer, pH 5.0, at a ratio of 2.5:1 (lipid:mRNA). The mixture was neutralized with Tris-HCl, pH 7.5, sucrose was added as a cryoprotectant, and the final solution was sterile-filtered. Vials were filled with formulated lipid nanoparticle and stored frozen at –20°C until further use. The pre-clinical vaccine product underwent analytical characterization, which included the determination of particle size and polydispersity, encapsulation, mRNA purity, double-stranded RNA content, osmolality, pH, endotoxin, and bioburden, and the material was deemed acceptable for *in vivo* study.

#### Viral antigens

Recombinant soluble S and RBD proteins from Wuhan-1 and B.1.1.529 SARS-CoV-2 strains were expressed as described (Amanat et al., 2021; Stadlbauer et al., 2020). Recombinant proteins were produced in Expi293F cells (ThermoFisher) by transfection of DNA using the ExpiFectamine 293 Transfection Kit (ThermoFisher). Supernatants were harvested 3 days post-transfection, and recombinant proteins were purified using Ni-NTA agarose (ThermoFisher), then buffer exchanged into PBS and concentrated using Amicon Ultracel centrifugal filters (EMD Millipore).

#### ELISA

Assays were performed in 96-well microtiter plates (Thermo Fisher) coated with 50 μL of recombinant Wuhan-1 or B.1.1.529 spike or RBD proteins. Plates were incubated at 4°C overnight and then blocked with 200 μL of 3% non-fat dry milk (AmericanBio) in PBS containing 0.1% Tween-20 (PBST) for 1 h at room temperature (RT). Sera were serially diluted in 1% non-fat dry milk in PBST and added to the plates. Plates were incubated for 2 h at room temperature and then washed 3 times with PBST. Goat anti-mouse IgG-HRP (Sigma-Aldrich, 1:9000) was diluted in 1% non-fat dry milk in PBST before adding to the wells and incubating for 1 h at room temperature. Plates were washed 3 times with PBST before the addition of peroxidase substrate (SigmaFAST o-phenylenediamine dihydrochloride, Sigma-Aldrich). Reactions were stopped by the addition of 3 M hydrochloric acid. Optical density (OD) measurements were taken at 490 nm, and endpoint titers were calculated in excel using a 0.15 OD 490 nm cutoff. Graphs were generated using Graphpad Prism v9.

#### Focus reduction neutralization test

Serial dilutions of sera were incubated with 10^2^ focus-forming units (FFU) of WA1/2020 D614G or B.1.1.529 for 1 h at 37°C. Antibody-virus complexes were added to Vero-TMPRSS2 cell monolayers in 96-well plates and incubated at 37°C for 1 h. Subsequently, cells were overlaid with 1% (w/v) methylcellulose in MEM. Plates were harvested 30 h later by removing overlays and fixed with 4% PFA in PBS for 20 min at room temperature. Plates were washed and sequentially incubated with an oligoclonal pool of SARS2-2, SARS2-11, SARS2-16, SARS2-31, SARS2-38, SARS2-57, and SARS2-71(Liu et al., 2021c) anti-S antibodies and HRP-conjugated goat anti-mouse IgG (Sigma Cat # A8924, RRID: AB_258426) in PBS supplemented with 0.1% saponin and 0.1% bovine serum albumin. SARS-CoV-2-infected cell foci were visualized using TrueBlue peroxidase substrate (KPL) and quantitated on an ImmunoSpot microanalyzer (Cellular Technologies).

#### Pseudovirus neutralization assay

Codon-optimized full-length spike genes (Wuhan-1 with D614G, B. 1.351, B. 1.617.2, and B.1.1.529) were cloned into a pCAGGS vector. Spike genes of contained the following mutations: B.1.351: L18F-D80A-D215G-L242-244del-R246I-K417N-E484K-N501Y-D614G-A701V; B.1.617.2: T19R-T95I-G142D-E156G-F157del-R158del-L452R-T478K-D614G-P681R-D950N; and B.1.1.529: A67V, Δ69-70, T95I, G142D/ΔVYY143-145, ΔN211/L212I, ins214EPE, G339D, S371L, S373P, S375F, K417N, N440K, G446S, S477N, T478K, E484A, Q493R, G496S, Q498R, N501Y, Y505H, T547K, D614G, H655Y, N679K, P681H, N764K, D796Y, N856K, Q954H, N969K, L981F. To generate VSVΔG-based SARS-CoV-2 pseudovirus, BHK-21/WI-2 cells were transfected with the spike expression plasmid and infected by VSVΔG-firefly-luciferase as previously described (Whitt, 2010). A549-hACE2-TMPRSS2 cells were used as target cells for the neutralization assay and maintained in DMEM supplemented with 10% fetal bovine serum and 1 μg/mL puromycin. To perform neutralization assay, mouse serum samples were heat-inactivated for 45 min at 56°C, and serial dilutions were made in DMEM supplemented with 10% FBS. The diluted serum samples or culture medium (serving as virus only control) were mixed with VSVΔG-based SARS-CoV-2 pseudovirus and incubated at 37°C for 45 min. The inoculum virus or virus-serum mix was subsequently used to infect A549-hACE2-TMPRSS2 cells for 18 h at 37°C. At 18 h post infection, an equal volume of One-Glo reagent (Promega; E6120) was added to culture medium for readout using BMG PHERastar-FS plate reader. The percentage of neutralization was calculated based on relative light units of the virus control, and subsequently analyzed using four parameter logistic curve (Prism 8,0).

#### Mouse experiments

##### (a) K18hACE2 transgenic mice

Seven-week-old female K18-hACE2 C57BL/6 mice were immunized three weeks apart with 5, 0.25, or 0.1 μg of mRNA vaccines (control or mRNA-1273) in 50 μl of PBS via intramuscular injection in the hind leg. Animals were bled three to four weeks after the second vaccine dose for immunogenicity analysis. In some experiments, four weeks after completing the primary series immunization, mice were challenged with 10^4^ FFU of WA1/2020 D614G or B.1.1.529 of SARS-CoV-2 strains by the intranasal route. In other experiments, 17 to 19 weeks after completing the primary series immunization, animals were bled for antibody analysis, and then boosted with 1 μg of control or mRNA-1273 vaccine. Four weeks later, K18-hACE2 mice were challenged with 10^4^ FFU of B.1.1.529 by the intranasal route. For some studies, weights were measured at day 0 and 6, and in all experiments, animals were euthanized at 6 dpi. Tissues were harvested for virological, immunological, and pathological analyses.

##### (b) BALB/c mice

6 to 8-week-old female BALB/c mice were immunized three weeks apart with 1 or 0.1 μg of mRNA vaccines (mRNA-1273 or mRNA-1273.529) or PBS (in 50 μL) via intramuscular injection in the quadriceps muscle of the hind leg under isoflurane anesthesia. Blood was sampled three weeks after the first immunization and two weeks after the second immunization, and anti-spike and neutralizing antibody levels were measured by ELISA and a VSV-based pseudovirus neutralization assay.

##### (c) 129S2 mice

Seven-week-old female 129S2 mice were immunized three weeks apart with 5 or 0.25 μg of mRNA vaccines (control, mRNA-1273) in 50 μl of PBS via intramuscular injection in the hind leg. Ten to eleven weeks later, animals were bled (pre-boost), and subsequently boosted with 1 μg of control, mRNA-1273, or mRNA-1273.529 vaccine. Three to four weeks later, animals were bled again (pos-boost), and then challenged with 10^4^ FFU of B.1.1.529 by the intranasal route. Animals were euthanized at 3 dpi, and tissues were harvested for virological and cytokine/chemokine analyses.

#### Measurement of viral burden

Tissues were weighed and homogenized with zirconia beads in a MagNA Lyser instrument (Roche Life Science) in 1 ml of DMEM medium supplemented with 2% heat-inactivated FBS. Tissue homogenates were clarified by centrifugation at 10,000 rpm for 5 min and stored at −80°C. RNA was extracted using the MagMax mirVana Total RNA isolation kit (Thermo Fisher Scientific) on the Kingfisher Flex extraction robot (Thermo Fisher Scientific). RNA was reverse transcribed and amplified using the TaqMan RNA-to-CT 1-Step Kit (Thermo Fisher Scientific). Reverse transcription was carried out at 48°C for 15 min followed by 2 min at 95°C. Amplification was accomplished over 50 cycles as follows: 95°C for 15 s and 60°C for 1 min. Copies of SARS-CoV-2 *N* gene RNA in samples were determined using a published assay (Case et al., 2020a).

#### Cytokine and chemokine protein measurements

Lung homogenates were incubated with Triton-X-100 (1% final concentration) for 1 h at room temperature to inactivate SARS-CoV-2. Homogenates were analyzed for cytokines and chemokines by Eve Technologies Corporation (Calgary, AB, Canada) using their Mouse Cytokine Array/Chemokine Array 31-Plex (MD31) platform.

#### Lung histology

Lungs of euthanized mice were inflated with ~2 mL of 10% neutral buffered formalin using a 3-mL syringe and catheter inserted into the trachea and kept in fixative for 7 days. Tissues were embedded in paraffin, and sections were stained with hematoxylin and eosin. Images were captured using the Nanozoomer (Hamamatsu) at the Alafi Neuroimaging Core at Washington University.

### QUANTIFICATION AND STATISTICAL ANALYSES

Statistical significance was assigned when *P* values were < 0.05 using GraphPad Prism version 9.3. Tests, number of animals, median values, and statistical comparison groups are indicated in the Figure legends. Changes in infectious virus titer, viral RNA levels, or serum antibody responses were compared to unvaccinated or mRNA-control immunized animals and were analyzed by one-way ANOVA with a multiple comparisons correction, Mann-Whitney test, or Wilcoxon signed-rank test depending on the type of results, number of comparisons, and distribution of the data.

