## Supplementary figures and images for "Boosting with Omicron-matched or historical mRNA vaccines increases neutralizing antibody responses and protection against B.1.1.529 infection in mice"

### Supplemental Figure S1

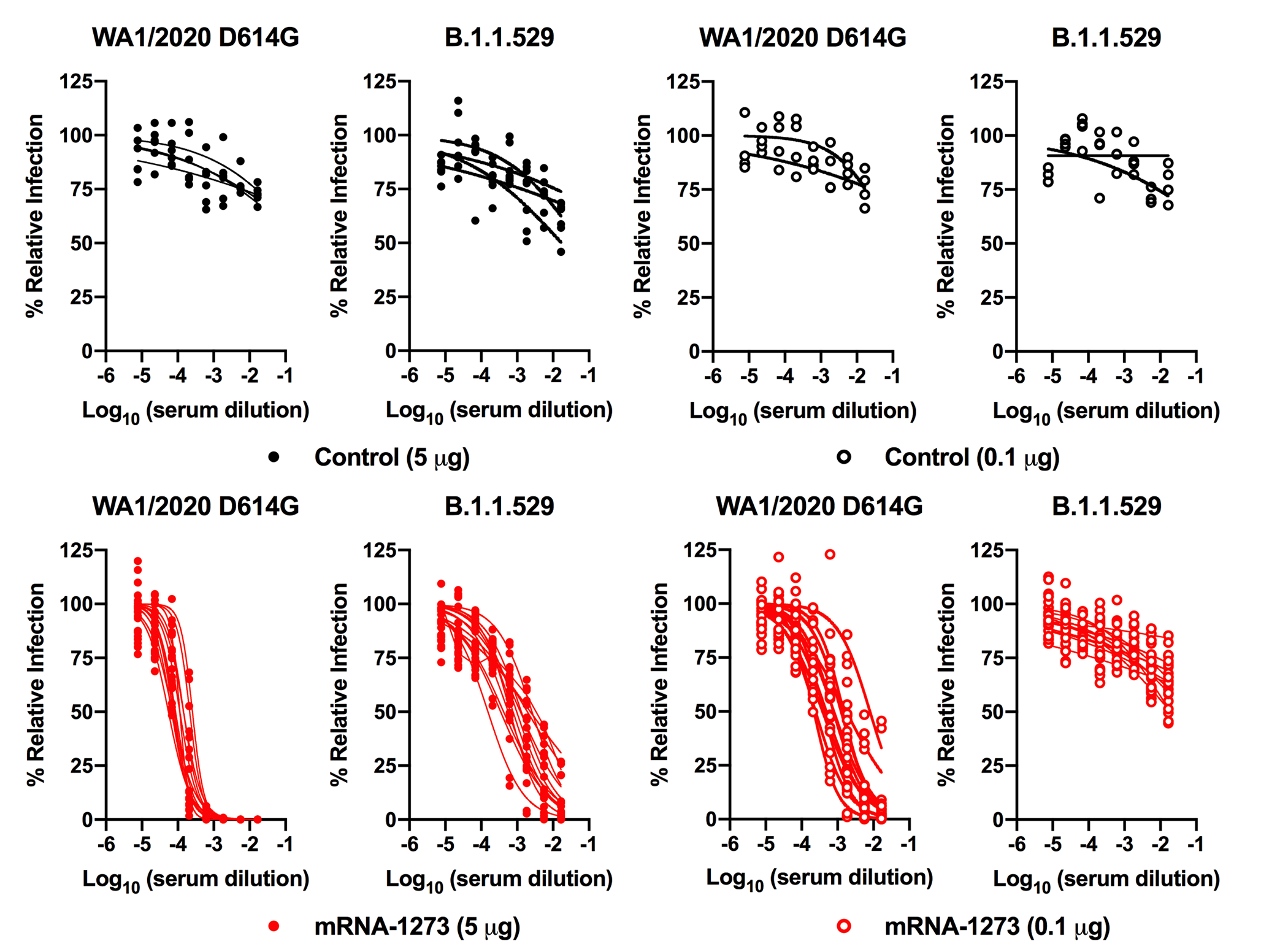

### Supplemental Figure S2

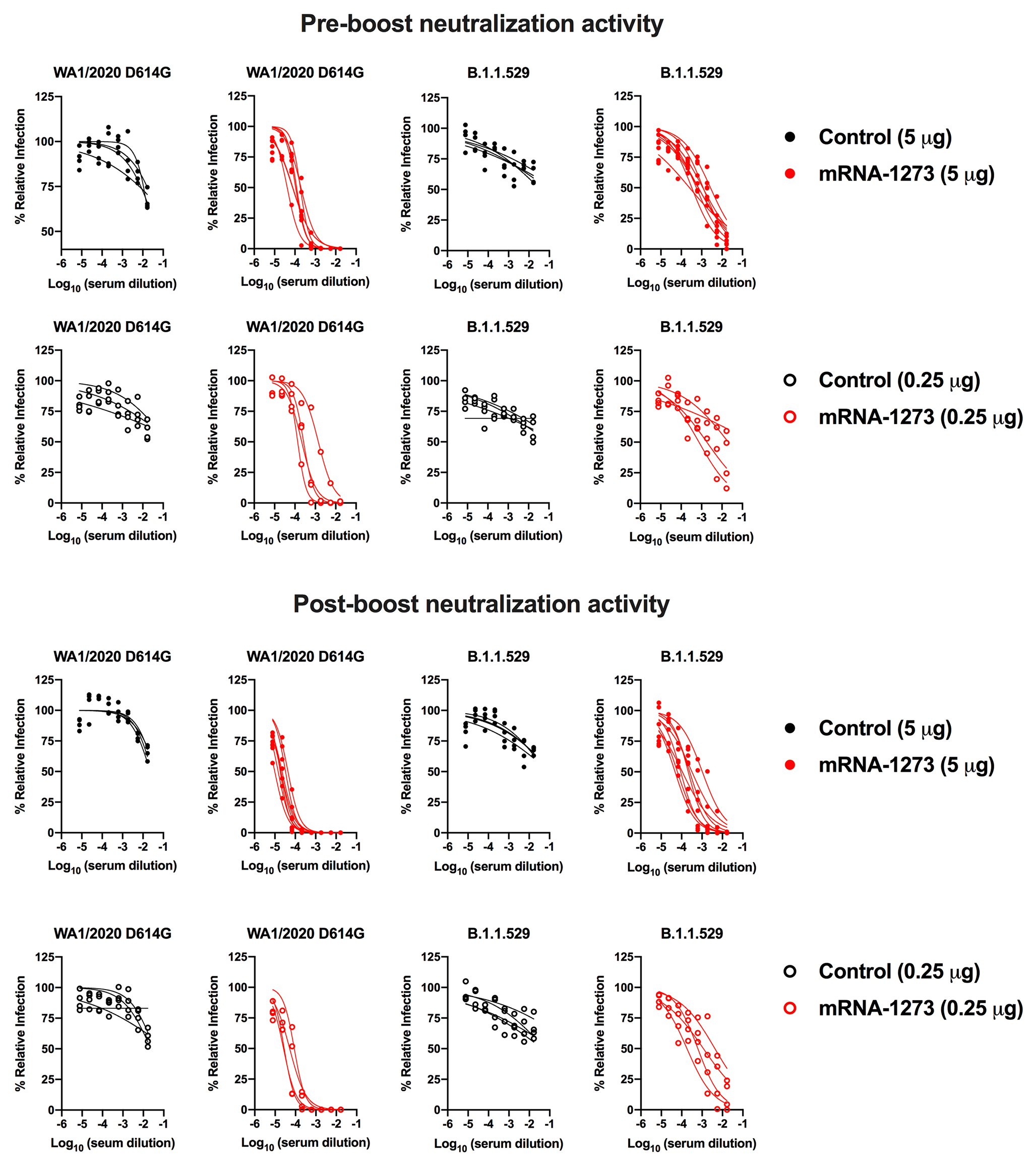

### Supplemental Figure S3

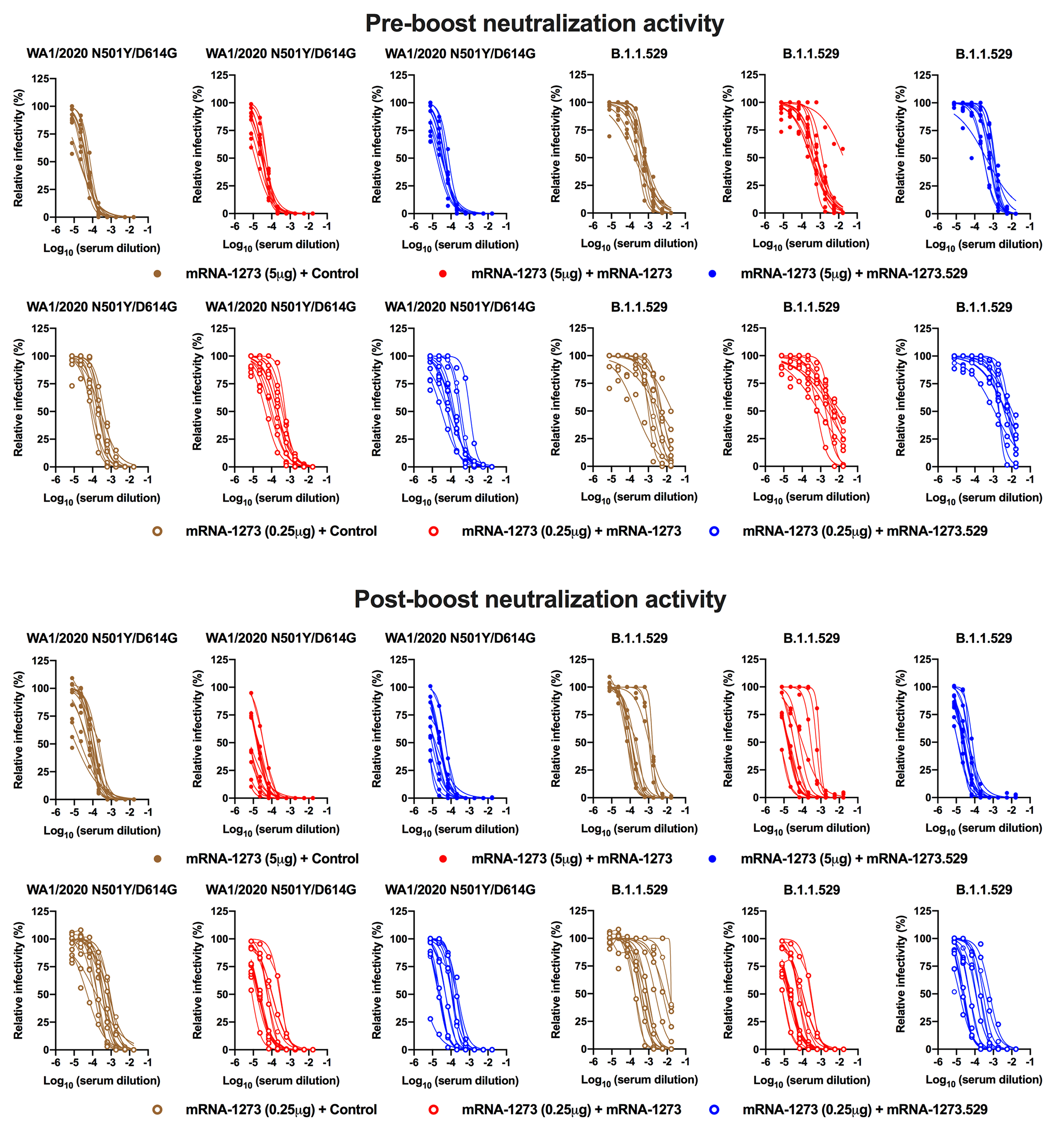
