## Supplementary Table S1 for "Boosting with Omicron-matched or historical mRNA vaccines increases neutralizing antibody responses and protection against B.1.1.529 infection in mice"

**Supplemental Table S1: Cytokine and chemokine concentrations in K18-hACE2 mice vaccinated with two 5 μg doses of mRNA vaccines and challenged with WA1/2020 D614G or B.1.1.529, Related to Figure 3.**

| **Cytokine/**  **Chemokine** | **Concentration (mean ± SD (pg/mL))** | | | | |
| --- | --- | --- | --- | --- | --- |
|  | **Naive** | **WA1/2020 D614G** | | **B.1.1.529** | |
|  |  | **Control** | **mRNA-1273** | **Control** | **mRNA-1273** |
| **Eotaxin** | 85.4 ± 32.8 | 301.9 ± 124.1 | 207.3 ± 42.6 | 244.9 ± 60.4 | 234.3 ± 69.4 |
| **G-CSF** | 0.6 ± 0.0 | 72.2 ± 45.6 | 2.6 ± 0.7 | 20.4 ± 24.3 | 2.3 ± 0.5 |
| **GM-CSF** | 0.6 ± 0.0 | 18.7 ± 8.9 | 5.4 ± 1.9 | 13.4 ± 4.5 | 5.1 ± 1.3 |
| **IFNγ** | 1.4 ± 0.7 | 85.4 ± 42.2 | 1.7 ± 0.8 | 32.0 ± 22.5 | 1.7 ± 0.7 |
| **IL-1α** | 8.9 ± 9.6 | 25.9 ± 10.0 | 21.2 ± 4.3 | 41.8 ± 8.8 | 22.6 ± 8.2 |
| **IL-1β** | 0.5 ± 0.0 | 6.5 ± 7.2 | 2.0 ± 0.5 | 5.8 ± 3.9 | 2.0 ± 0.9 |
| **IL-2** | 4.3 ± 1.4 | 20.5 ± 5.1 | 6.3 ± 1.1 | 15.2 ± 6.3 | 6.6 ± 1.2 |
| **IL-3** | 0.6 ± 0.0 | 5.6 ± 2.6 | 0.9 ± 0.2 | 5.6 ± 5.1 | 0.9 ± 0.1 |
| **IL-4** | 0.6 ± 0.0 | 0.6 ± 0.0 | 0.6 ± 0.0 | 1.4 ± 0.9 | 0.7 ± 0.2 |
| **IL-5** | 0.6 ± 0.0 | 3.3 ± 2.2 | 0.9 ± 0.2 | 5.0 ± 3.8 | 1.1 ± 0.8 |
| **IL-6** | 1.1 ± 0.0 | 114.7 ± 137.6 | 2.0 ± 0.9 | 133.5 ± 118.0 | 2.2 ± 1.0 |
| **IL-7** | 0.7 ± 0.0 | 3.6 ± 0.8 | 3.9 ± 1.0 | 3.6 ± 1.2 | 2.9 ± 1.2 |
| **IL-9** | 34.3 ± 14.4 | 50.6 ± 12.2 | 55.6 ± 14.3 | 42.2 ± 11.8 | 61.9 ± 11.0 |
| **IL-10** | 0.6 ± 0.0 | 8.2 ± 5.1 | 2.9 ± 0.6 | 5.3 ± 1.7 | 2.7 ± 0.5 |
| **IL-12p70** | 1.5 ± 1.0 | 3.6 ± 1.0 | 3.1 ± 0.8 | 3.5 ± 1.0 | 2.6 ± 0.6 |
| **IL-15** | 5.0 ± 0.0 | 21.0 ± 3.3 | 14.0 ± 3.9 | 20.4 ± 4.9 | 13.0 ± 3.5 |
| **IL-17** | 0.6 ± 0.0 | 1.0 ± 0.5 | 0.6 ± 0.0 | 1.1 ± 0.7 | 0.6 ± 0.0 |
| **IP-10 (CXCL10)** | 21.0 ± 5.6 | 6616.0 ± 4061.4 | 55.8 ± 60.2 | 4693.8 ± 2493.1 | 42.2 ± 16.7 |
| **KC (CXCL1)** | 19.2 ± 5.2 | 129.2 ± 54.9 | 44.9 ± 31.6 | 161.4 ± 108.2 | 34.1 ± 16.8 |
| **LIF** | 0.9 ± 0.1 | 45.4 ± 40.8 | 1.6 ± 0.3 | 12.3 ± 8.2 | 1.5 ± 0.7 |
| **LIX (CXCL5)** | 23.4 ± 0.0 | 1078.1 ± 1061.8 | 964.7 ± 639.8 | 1697.2 ± 1236.9 | 1143.8 ± 1040.2 |
| **MCP-1 (CCL2)** | 22.6 ± 6.1 | 1587.6 ± 1155.2 | 33.9 ± 22.7 | 605.8 ± 439.5 | 22.0 ± 13.3 |
| **M-CSF** | 5.0 ± 1.7 | 13.1 ± 5.5 | 7.7 ± 1.3 | 12.3 ± 4.8 | 6.7 ± 2.0 |
| **MIG (CXCL9)** | 42.3 ± 20.7 | 10000.0 ± 0 | 131.7 ± 124.5 | 9270.7 ± 2062.8 | 120.0 ± 85.0 |
| **MIP-1α (CCL3)** | 32.9 ± 20.7 | 879.6 ± 507.6 | 36.3 ± 43.9 | 171.0 ± 103.4 | 20.5 ± 18.7 |
| **MIP-1β (CCL4)** | 9.0 ± 0.0 | 622.9 ± 426.0 | 21.5 ± 10.0 | 122.7 ± 59.1 | 18.4 ± 4.9 |
| **MIP-2 (CXCL2)** | 42.9 ± 5.4 | 112.6 ± 31.5 | 92.7 ± 9.0 | 136.0 ± 27.8 | 105.3 ± 19.4 |
| **RANTES (CCL5)** | 57.2 ± 19.3 | 110.5 ± 50.1 | 41.5 ± 23.9 | 62.6 ± 22.6 | 35.8 ± 13.2 |
| **TNFα** | 0.6 ± 0.0 | 12.5 ± 47.0 | 1.7 ± 0.9 | 5.8 ± 2.1 | 1.2 ± 0.5 |

Cytokine and chemokine levels in lung homogenates at 6 dpi in animals immunized with two 5 μg doses of indicated mRNA vaccines. Data are expressed as mean ± standard deviation in pg/mL (2 experiments, n = 7-8 per group except naive, n = 4).
