## Supplementary Table S2 for "Boosting with Omicron-matched or historical mRNA vaccines increases neutralizing antibody responses and protection against B.1.1.529 infection in mice"

**Supplemental Table S2: Cytokine and chemokine concentrations in K18-hACE2 mice vaccinated with two 0.1 μg doses of mRNA vaccines and challenged with WA1/2020 D614G or B.1.1.529, Related to Figure 3.**

| **Cytokine/**  **Chemokine** | **Concentration (mean ± SD (pg/mL))** | | | | |
| --- | --- | --- | --- | --- | --- |
|  | **Naive** | **WA1/2020 D614G** | | **B.1.1.529** | |
|  |  | **Control** | **mRNA-1273** | **Control** | **mRNA-1273** |
| **Eotaxin** | 85.4 ± 32.8 | 322.0 ± 90.1 | 263.6 ± 62.4 | 204.3 ± 27.7 | 711.0 ± 287.6 |
| **G-CSF** | 0.6 ± 0.0 | 57.2 ± 13.6 | 3.2 ± 2.1 | 25.9 ± 22.2 | 29.8 ± 18.8 |
| **GM-CSF** | 0.6 ± 0.0 | 19.8 ± 8.0 | 6.3 ± 2.0 | 16.4 ± 4.2 | 25.5 ± 10.6 |
| **IFNγ** | 1.4 ± 0.7 | 75.7 ± 36.7 | 1.7 ± 0.8 | 29.3 ± 17.1 | 15.6 ± 15.9 |
| **IL-1α** | 8.9 ± 9.6 | 23.3 ± 5.8 | 19.4 ± 5.4 | 32.0 ± 7.3 | 34.3 ± 10.2 |
| **IL-1β** | 0.5 ± 0.0 | 4.7 ± 2.7 | 3.1 ± 3.6 | 7.8 ± 7.4 | 17.5 ± 6.6 |
| **IL-2** | 4.3 ± 1.4 | 17.5 ± 6.6 | 6.2 ± 1.3 | 17.9 ± 4.7 | 8.4 ± 2.6 |
| **IL-3** | 0.6 ± 0.0 | 5.1 ± 2.5 | 0.8 ± 0.3 | 6.5 ± 4.7 | 2.1 ± 0.8 |
| **IL-4** | 0.6 ± 0.0 | 0.6 ± 0.0 | 0.7 ± 0.2 | 1.9 ± 1.3 | 13.4 ± 8.2 |
| **IL-5** | 0.6 ± 0.0 | 4.6 ± 3.1 | 3.0 ± 1.7 | 6.5 ± 4.9 | 133.5 ± 110.0 |
| **IL-6** | 1.1 ± 0.0 | 82.7 ± 61.4 | 4.0 ± 4.0 | 161.4 ± 133.2 | 329.7 ± 237.1 |
| **IL-7** | 0.7 ± 0.0 | 3.1 ± 0.6 | 3.3 ± 0.9 | 3.4 ± 0.6 | 3.9 ± 0.8 |
| **IL-9** | 34.3 ± 14.4 | 56.2 ± 13.0 | 45.1 ± 19.1 | 51.0 ± 16.0 | 61.0 ± 12.4 |
| **IL-10** | 0.6 ± 0.0 | 8.6 ± 3.6 | 2.7 ± 0.7 | 4.5 ± 1.1 | 4.2 ± 1.0 |
| **IL-12p70** | 1.5 ± 1.0 | 3.7 ± 0.4 | 2.7 ± 1.0 | 2.8 ± 1.0 | 2.9 ± 0.7 |
| **IL-15** | 5.0 ± 0.0 | 18.9 ± 3.6 | 15.0 ± 5.1 | 16.8 ± 5.1 | 17.5 ± 4.0 |
| **IL-17** | 0.6 ± 0.0 | 0.8 ± 0.3 | 0.6 ± 0.0 | 1.2 ± 0.8 | 0.8 ± 0.2 |
| **IP-10 (CXCL10)** | 21.0 ± 5.6 | 6211.1 ± 3501.0 | 86.0 ± 77.0 | 4915.5 ± 2589.8 | 1571.3 ± 1414.3 |
| **KC (CXCL1)** | 19.2 ± 5.2 | 177.0 ± 145.8 | 51.3 ± 30.2 | 164.7 ± 76.8 | 362.7 ± 284.7 |
| **LIF** | 0.9 ± 0.1 | 40.9 ± 17.1 | 1.9 ± 0.8 | 15.8 ± 7.9 | 20.4 ± 8.4 |
| **LIX (CXCL5)** | 23.4 ± 0.0 | 501.4 ± 294.7 | 1377.9 ± 778.3 | 1207.5 ± 967.9 | 289.3 ± 180.2 |
| **MCP-1 (CCL2)** | 22.6 ± 6.1 | 1360.2 ± 835.5 | 47.5 ± 29.4 | 567.3 ± 294.7 | 554.6 ± 298.1 |
| **M-CSF** | 5.0 ± 1.7 | 13.5 ± 3.8 | 9.2 ± 4.8 | 17.1 ± 5.5 | 19.1 ± 5.7 |
| **MIG (CXCL9)** | 42.3 ± 20.7 | 8773.3 ± 3469.5 | 299.1 ± 339.4 | 9007.5 ± 2625.8 | 8885.3 ± 2800.0 |
| **MIP-1α (CCL3)** | 32.9 ± 20.7 | 655.9 ± 294.6 | 35.1 ± 30.3 | 168.3 ± 107.6 | 203.4 ± 161.6 |
| **MIP-1β (CCL4)** | 9.0 ± 0.0 | 435.2 ± 191.3 | 28.2 ± 15.5 | 118.0 ± 86.6 | 142.7 ±100.4 |
| **MIP-2 (CXCL2)** | 42.9 ± 5.4 | 100.0 ± 15.7 | 106.0 ± 24.7 | 128.5 ± 32.8 | 234.3 ± 122.8 |
| **RANTES (CCL5)** | 57.2 ± 19.3 | 112.2 ± 44.4 | 56.4 ± 38.7 | 67.7 ± 31.6 | 104.1 ± 28.0 |
| **TNFα** | 0.6 ± 0.0 | 10.3 ± 3.9 | 1.7 ± 0.8 | 6.4 ± 2.7 | 9.0 ± 2.6 |

Cytokine and chemokine levels in lung homogenates at 6 dpi in animals immunized with two 0.1 μg doses of indicated mRNA vaccines. Data are expressed as mean ± standard deviation in pg/mL (2 experiments, n = 7-8 per group except naive, n = 4).
