## Supplementary Table S3 for "Boosting with Omicron-matched or historical mRNA vaccines increases neutralizing antibody responses and protection against B.1.1.529 infection in mice"

**Supplemental Table S3: Cytokine and chemokine concentrations in 129S2 mice immunized with 5 µg primary series and 1 µg booster dose and challenged with WA1/2020 N501Y/D614G or B.1.1.529, Related to Figure 7.**

| **Challenge virus** |  | **WA1/2020 N501Y/D614G** | **WA1/2020 N501Y/D614G** | **WA1/2020 N501Y/D614G** | **WA1/2020 N501Y/D614G** | **B.1.1.529** | **B.1.1.529** | **B.1.1.529** | **B.1.1.529** |
| --- | --- | --- | --- | --- | --- | --- | --- | --- | --- |
| **Vaccine sequence** | **Naive** | **Control (1)**  **Control (2)**  **Control (3)** | **mRNA-1273 (1)**  **mRNA-1273 (2)**  **Control (3)** | **mRNA-1273 (1)**  **mRNA-1273 (2)**  **mRNA-1273 (3)** | **mRNA-1273 (1)**  **mRNA-1273 (2)**  **mRNA-1273.529 (3)** | **Control (1)**  **Control (2)**  **Control (3)** | **mRNA-1273 (1)**  **mRNA-1273 (2)**  **Control (3)** | **mRNA-1273 (1)**  **mRNA-1273 (2)**  **mRNA-1273 (3)** | **mRNA-1273 (1)**  **mRNA-1273 (2)**  **mRNA-1273.529 (3)** |
| **Cytokine/**  **Chemokine** | **Concentration (mean ± SD (pg/mL))** | | | | | | | | |
| **Eotaxin** | **222.5 ± 55.5** | **139.5 ± 22.4** | **91.1 ± 33.8** | **218.8 ± 144.2** | **143.2 ± 20.1** | **95.2 ± 35.8** | **133.3 ± 44.8** | **137.3 ± 36.9** | **89.1 ± 12.8** |
| **G-CSF** | **2 ± 0.6** | **348.9 ± 163.5** | **3.1 ± 0.4** | **8.6 ± 11** | **4 ± 0.3** | **9.4 ± 4.4** | **4 ± 1** | **2.8 ± 0.8** | **2.1 ± 0.8** |
| **GM-CSF** | **7.2 ± 0.9** | **24.9 ± 8** | **6.4 ± 1.3** | **9 ± 3.4** | **8.5 ± 1.2** | **9.8 ± 0.8** | **6 ± 1.1** | **8 ± 0.6** | **5.2 ± 1.8** |
| **IFN-γ** | **2.1 ± 0.7** | **3 ± 0.8** | **1.2 ± 0.9** | **1.1 ± 0.7** | **2.2 ± 0.6** | **3 ± 1.1** | **1.6 ± 1.2** | **1 ± 0.3** | **1.2 ± 0.7** |
| **IL-1-α** | **27.2 ± 14** | **201.1 ± 135.2** | **19.7 ± 7.3** | **54.3 ± 86.4** | **28.3 ± 10.4** | **52.4 ± 20.9** | **20.1 ± 27.4** | **19.7 ± 12.4** | **24.2 ± 22** |
| **IL-1-β** | **1.4 ± 0.6** | **17.5 ± 9** | **2.2 ± 0.8** | **3.4 ± 3** | **2.8 ± 0.4** | **3 ± 0.6** | **1.8 ± 0.4** | **1.3 ± 0.5** | **1.6 ± 0.7** |
| **IL-2** | **7.8 ± 1.5** | **6.7 ± 0.8** | **7.8 ± 2.1** | **5.7 ± 1** | **7.4 ± 1.3** | **10.5 ± 3.1** | **8.9 ± 5.3** | **6.7 ± 1.6** | **9.4 ± 5.8** |
| **IL-3** | **1.1 ± 0.2** | **1.4 ± 0.2** | **0.9 ± 0.3** | **0.9 ± 0.1** | **1.1 ± 0.2** | **1.4 ± 0.2** | **0.8 ± 0.2** | **0.9 ± 0.2** | **0.8 ± 0.2** |
| **IL-4** | **0.7 ± 0** | **1 ± 0.6** | **0.8 ± 0.2** | **4.5 ± 7.6** | **1 ± 0.7** | **1 ± 0.5** | **0.7 ± 0** | **0.7 ± 0** | **0.7 ± 0.1** |
| **IL-5** | **0.8 ± 0.1** | **3.2 ± 1** | **2.8 ± 2** | **112 ± 183.5** | **3.2 ± 4.7** | **7.8 ± 9.2** | **1.2 ± 0.5** | **1.1 ± 0.4** | **1.6 ± 0.7** |
| **IL-6** | **1.6 ± 0.9** | **1395.6 ± 1007.1** | **1.9 ± 0.8** | **8.1 ± 13.6** | **1.9 ± 0.9** | **69.3 ± 57.7** | **3.5 ± 2.8** | **1.8 ± 0.8** | **1.3 ± 0.6** |
| **IL-7** | **4.6 ± 0.4** | **4 ± 0.5** | **2.6 ± 0.7** | **3.8 ± 0.5** | **4.3 ± 0.9** | **4.5 ± 0.8** | **2.9 ± 0.1** | **3.5 ± 0.3** | **2.7 ± 0.8** |
| **IL-9** | **41.1 ± 16.8** | **32 ± 2.9** | **23.8 ± 5.3** | **43.4 ± 15.1** | **35 ± 12.8** | **34.9 ± 12.1** | **33.1 ± 9.4** | **38.9 ± 4.2** | **27.5 ± 10.6** |
| **IL-10** | **3.8 ± 0.6** | **8.2 ± 4.7** | **3.4 ± 1.1** | **3.8 ± 0.4** | **4.7 ± 0.8** | **6.2 ± 1.6** | **3.4 ± 0.6** | **3.6 ± 0.8** | **3.6 ± 1.3** |
| **IL-12p70** | **2.7 ± 0.6** | **8 ± 6** | **2 ± 0.4** | **2.5 ± 0.3** | **3.1 ± 1** | **4.1 ± 2.1** | **2 ± 1** | **2.3 ± 0.4** | **2.3 ± 1.9** |
| **IL-15** | **16.9 ± 2.7** | **18.4 ± 2.3** | **20 ± 21** | **14.4 ± 0.5** | **18 ± 2.2** | **18.9 ± 2.5** | **9.6 ± 2.3** | **13.7 ± 1.4** | **8.6 ± 3.2** |
| **IP-10 (CXCL10)** | **15.5 ± 1.2** | **4703.3 ± 1807.1** | **26.4 ± 27.2** | **36.4 ± 51.5** | **14.1 ± 4.1** | **706.4 ± 613.5** | **103.6 ± 83.8** | **23 ± 11.6** | **12.6 ± 4.6** |
| **KC (CXCL1)** | **74.2 ± 27.5** | **1988.7 ± 859.3** | **70.9 ± 30.6** | **192.8 ± 197.6** | **116.9 ± 49.3** | **233.9 ± 101.6** | **163.2 ± 64** | **96 ± 29.2** | **74.5 ± 48.5** |
| **LIF** | **1.4 ± 0.2** | **14.8 ± 3.8** | **1.2 ± 0.4** | **3.7 ± 3.3** | **1.6 ± 0.1** | **2.3 ± 0.7** | **1.9 ± 0.8** | **1.5 ± 0.3** | **1.1 ± 0.4** |
| **LIX (CXCL5)** | **61.7 ± 17** | **111.1 ± 33.2** | **78.4 ± 50.1** | **102.9 ± 69.6** | **233.7 ± 243.1** | **69.2 ± 19.1** | **58.8 ± 20.9** | **183 ± 132.6** | **53.7 ± 19.9** |
| **MCP-1 (CCL2)** | **7.9 ± 1.4** | **2189.7 ± 732.3** | **12.1 ± 3.6** | **105.4 ± 187.7** | **11.4 ± 1.6** | **161.8 ± 138.3** | **44.2 ± 28.7** | **13.5 ± 3.9** | **8.1 ± 3.4** |
| **M-CSF** | **16 ± 4** | **27.8 ± 8.6** | **7.3 ± 3.1** | **23 ± 26.1** | **12.5 ± 2.7** | **17.3 ± 8.2** | **10.2 ± 3.8** | **9.5 ± 1.7** | **10.3 ± 4.6** |
| **MIG (CXCL9)** | **23.4 ± 15.2** | **847.2 ± 308.9** | **60.5 ± 91.7** | **21.2 ± 5.5** | **22.7 ± 5.3** | **174.1 ± 168.9** | **251.9 ± 167** | **90 ± 52.6** | **30 ± 13.3** |
| **MIP-1 (CCL3)** | **44.3 ± 32.8** | **353.4 ± 105.4** | **243.3 ± 81.4** | **165.7 ± 147.3** | **335.6 ± 159.7** | **409.5 ± 147.8** | **279.8 ± 257.2** | **244.5 ± 184.3** | **284.7 ± 211.6** |
| **MIP-1 (CCL4)** | **16 ± 3.5** | **146.7 ± 58.9** | **14.7 ± 2.8** | **36.4 ± 42.7** | **21.4 ± 1.5** | **33.9 ± 9.7** | **15.2 ± 5.8** | **14.8 ± 5** | **11.6 ± 1.9** |
| **MIP-2 (CXCL2)** | **121.8 ± 7.6** | **832 ± 279.6** | **89.4 ± 15.2** | **333 ± 478.4** | **113.3 ± 7.9** | **161.7 ± 13.4** | **84.4 ± 23.4** | **98.4 ± 4.8** | **93.9 ± 19.8** |
| **RANTES (CCL5)** | **15.5 ± 3.3** | **35.1 ± 7.5** | **11.9 ± 6.2** | **45 ± 11.8** | **48.8 ± 5** | **20.5 ± 10.7** | **24.8 ± 13.4** | **64.3 ± 11.1** | **20.7 ± 6.5** |
| **TNF-α** | **0.7 ± 0** | **11.1 ± 4.7** | **1.7 ± 1.2** | **1.8 ± 1.4** | **3.6 ± 2.5** | **7.5 ± 3.5** | **4.9 ± 8** | **2.6 ± 2.6** | **6.3 ± 8.3** |

Cytokine and chemokine levels in lung homogenates at 3 dpi with WA1/2020 D614G or B.1.1.529 in animals immunized with a primary series of two 5 μg doses of control or mRNA-1273 vaccines (labeled 1 and 2) and then boosted with control, mRNA-1273, or mRNA-1273.529 vaccines (labeled 3). Data are expressed as mean ± standard deviation in pg/mL (2 experiments, n = 8 per group except naive, n = 4).
