## Supplementary Table S4 for "Boosting with Omicron-matched or historical mRNA vaccines increases neutralizing antibody responses and protection against B.1.1.529 infection in mice"

**Supplemental Table S4: Cytokine and chemokine concentrations in 129S2 mice immunized with 0.25 µg primary series and 1 µg booster dose and challenged with WA1/2020 N501Y/D614G or B.1.1.529, Related to Figure 7.**

| **Challenge virus** |  | **WA1/2020 N501Y/D614G** | **WA1/2020 N501Y/D614G** | **WA1/2020 N501Y/D614G** | **WA1/2020 N501Y/D614G** | **B.1.1.529** | **B.1.1.529** | **B.1.1.529** | **B.1.1.529** |
| --- | --- | --- | --- | --- | --- | --- | --- | --- | --- |
| **Vaccine sequence** | **Naive** | **Control (1)**  **Control (2)**  **Control (3)** | **mRNA-1273 (1)**  **mRNA-1273 (2)**  **Control (3)** | **mRNA-1273 (1)**  **mRNA-1273 (2)**  **mRNA-1273 (3)** | **mRNA-1273 (1)**  **mRNA-1273 (2)**  **mRNA-1273.529 (3)** | **Control (1)**  **Control (2)**  **Control (3)** | **mRNA-1273 (1)**  **mRNA-1273 (2)**  **Control (3)** | **mRNA-1273 (1)**  **mRNA-1273 (2)**  **mRNA-1273 (3)** | **mRNA-1273 (1)**  **mRNA-1273 (2)**  **mRNA-1273.529 (3)** |
| **Cytokine/**  **Chemokine** | **Concentration (mean ± SD (pg/mL))** | | | | | | | | |
| **Eotaxin** | **222.5 ± 55.5** | **214.5 ± 79.1** | **164.7 ± 71.8** | **178.7 ± 98.5** | **140.7 ± 54.3** | **190.4 ± 43.8** | **441.2 ± 207.5** | **345.9 ± 89.5** | **193.9 ± 123.5** |
| **G-CSF** | **2 ± 0.6** | **192.4 ± 124.2** | **5.6 ± 1.6** | **5.3 ± 2.8** | **5.3 ± 0.9** | **13.6 ± 12.2** | **57.6 ± 69.3** | **6.3 ± 2.5** | **4.3 ± 1.1** |
| **GM-CSF** | **7.2 ± 0.9** | **29.3 ± 19.7** | **10.2 ± 1** | **9.8 ± 1.8** | **9.2 ± 2.6** | **12.3 ± 2.5** | **19.5 ± 11.3** | **11 ± 2.2** | **9.1 ± 1.6** |
| **IFN-γ** | **2.1 ± 0.7** | **4.1 ± 1.2** | **2.6 ± 0.8** | **1.8 ± 0.7** | **1.9 ± 0.6** | **2.6 ± 1.1** | **4.4 ± 1.9** | **1.3 ± 0.1** | **1.7 ± 0.1** |
| **IL-1-α** | **27.2 ± 14** | **200.4 ± 57.9** | **53.5 ± 9.5** | **50.1 ± 45.8** | **52.1 ± 36.3** | **60.8 ± 18.8** | **75.4 ± 35.9** | **59.4 ± 52.7** | **51.1 ± 39.2** |
| **IL-1-β** | **1.4 ± 0.6** | **16.1 ± 4.2** | **3.7 ± 0.9** | **4 ± 1.9** | **3.4 ± 1** | **5.1 ± 2.4** | **7 ± 3.1** | **3.1 ± 1.1** | **3.6 ± 0.3** |
| **IL-2** | **7.8 ± 1.5** | **5.2 ± 1** | **11.7 ± 3.6** | **9.7 ± 3.8** | **11.5 ± 3.5** | **7.9 ± 1.5** | **11.1 ± 2.1** | **7.6 ± 3.3** | **6.9 ± 1.6** |
| **IL-3** | **1.1 ± 0.2** | **1.5 ± 0.4** | **1.2 ± 0.2** | **1 ± 0.2** | **1 ± 0.2** | **1.4 ± 0.2** | **2.7 ± 1.3** | **1 ± 0.4** | **1.1 ± 0.3** |
| **IL-4** | **0.7 ± 0** | **0.7 ± 0** | **1.6 ± 1** | **1.3 ± 1.1** | **1 ± 0.5** | **1.6 ± 1.1** | **3.3 ± 2.1** | **2.8 ± 1.4** | **1.6 ± 1.1** |
| **IL-5** | **0.8 ± 0.1** | **3.1 ± 2.3** | **6.9 ± 5.6** | **30.1 ± 53.2** | **6.6 ± 6.6** | **14.4 ± 18.6** | **62.7 ± 40.6** | **22.7 ± 23.2** | **8.8 ± 9.1** |
| **IL-6** | **1.6 ± 0.9** | **821.4 ± 528.8** | **4.3 ± 3** | **4.3 ± 3.1** | **2.3 ± 0.8** | **71.6 ± 79.8** | **165.1 ± 212.3** | **7.2 ± 4.9** | **4.5 ± 3.2** |
| **IL-7** | **4.6 ± 0.4** | **4.2 ± 1.1** | **4.5 ± 0.5** | **4.6 ± 0.6** | **3.4 ± 0.8** | **5.3 ± 0.9** | **4.3 ± 1** | **4.6 ± 0.6** | **4.2 ± 1** |
| **IL-9** | **41.1 ± 16.8** | **39.8 ± 13.8** | **29.2 ± 5.1** | **34.5 ± 11.4** | **25.2 ± 9.5** | **45 ± 8.6** | **27.4 ± 5.2** | **27.3 ± 12** | **41 ± 11.2** |
| **IL-10** | **3.8 ± 0.6** | **6.4 ± 1** | **5.6 ± 1.3** | **4.2 ± 0.4** | **4.8 ± 0.6** | **6.2 ± 1.4** | **5.1 ± 1.5** | **5 ± 2.1** | **5.1 ± 0.6** |
| **IL-12p70** | **2.7 ± 0.6** | **5.2 ± 0.8** | **3.1 ± 0.9** | **2.8 ± 0.3** | **3.9 ± 1.6** | **4.2 ± 0.8** | **4.1 ± 1.1** | **6 ± 5.2** | **2.9 ± 0.4** |
| **IL-15** | **16.9 ± 2.7** | **21.9 ± 5.9** | **18.3 ± 4.6** | **17.2 ± 1.9** | **11.5 ± 2.9** | **21.4 ± 2.8** | **19.2 ± 4** | **15.5 ± 4.2** | **15.7 ± 2** |
| **IP-10 (CXCL10)** | **15.5 ± 1.2** | **4746.6 ± 1080.3** | **36.1 ± 42.6** | **27.5 ± 21.2** | **19.4 ± 6.7** | **1341.9 ± 855.1** | **2676.3 ± 3651** | **76.2 ± 61.9** | **21.7 ± 13.3** |
| **KC (CXCL1)** | **74.2 ± 27.5** | **1816.5 ± 1911.3** | **239.4 ± 92.8** | **225.1 ± 146.9** | **265.4 ± 201.5** | **373.6 ± 225.5** | **696.6 ± 656.3** | **275.6 ± 131** | **199.6 ± 149.7** |
| **LIF** | **1.4 ± 0.2** | **11.2 ± 3.4** | **1.9 ± 0.8** | **1.9 ± 0.5** | **1.5 ± 0.5** | **3.7 ± 1.1** | **8.1 ± 4.5** | **3.5 ± 1.2** | **2.2 ± 0.8** |
| **LIX (CXCL5)** | **61.7 ± 17** | **91.6 ± 58.8** | **218.2 ± 165** | **72 ± 17** | **68.4 ± 25.5** | **207.3 ± 126.6** | **267.2 ± 140.1** | **223.7 ± 54.9** | **93.1 ± 96.4** |
| **MCP-1 (CCL2)** | **7.9 ± 1.4** | **1639.5 ± 444.6** | **19.3 ± 9.1** | **19.6 ± 7.2** | **17.5 ± 7** | **286.6 ± 211.3** | **691.5 ± 839.4** | **45.7 ± 30.7** | **14.9 ± 7.1** |
| **M-CSF** | **16 ± 4** | **24.3 ± 7.6** | **15.1 ± 3.2** | **14.4 ± 4.4** | **14 ± 4.9** | **19.2 ± 4.7** | **22.9 ± 6.6** | **16.4 ± 1.8** | **11 ± 3.4** |
| **MIG (CXCL9)** | **23.4 ± 15.2** | **1139.2 ± 646.7** | **43.6 ± 33** | **37.1 ± 20.4** | **17.4 ± 2.5** | **303.1 ± 60.5** | **653 ± 561.5** | **76.4 ± 56.7** | **73.9 ± 15.8** |
| **MIP-1 (CCL3)** | **44.3 ± 32.8** | **310.1 ± 94.5** | **469.8 ± 92.2** | **341 ± 175.6** | **395.2 ± 164.9** | **372.9 ± 112.5** | **465.7 ± 161.3** | **199.2 ± 151.4** | **334.2 ± 29.8** |
| **MIP-1 (CCL4)** | **16 ± 3.5** | **139.1 ± 33.8** | **21.1 ± 3.5** | **25 ± 13.3** | **16 ± 4.1** | **41.7 ± 20.8** | **101.9 ± 95.7** | **23.7 ± 8.3** | **19.9 ± 2.2** |
| **MIP-2 (CXCL2)** | **121.8 ± 7.6** | **1022.7 ± 485.1** | **143.3 ± 26.3** | **242.8 ± 238.2** | **141.3 ± 62.3** | **263.8 ± 196.7** | **410.2 ± 277.2** | **207.9 ± 90.1** | **193.9 ± 133.2** |
| **RANTES (CCL5)** | **15.5 ± 3.3** | **51.1 ± 19.1** | **8.6 ± 1.1** | **23.8 ± 6.4** | **15.7 ± 6.1** | **20.4 ± 9** | **23.3 ± 16.1** | **24.8 ± 10.4** | **42.6 ± 23.3** |
| **TNF-α** | **0.7 ± 0** | **7.9 ± 1.2** | **6.2 ± 2.8** | **3 ± 1.7** | **7.7 ± 4.9** | **5.8 ± 2.4** | **7.6 ± 2.8** | **3.4 ± 4.1** | **3.6 ± 1.7** |

Cytokine and chemokine levels in lung homogenates at 3 dpi with WA1/2020 D614G or B.1.1.529 in animals immunized with a primary series of two 0.25 μg doses of control or mRNA-1273 vaccines (labeled 1 and 2) and then boosted with control, mRNA-1273, or mRNA-1273.529 vaccines (labeled 3). Data are expressed as mean ± standard deviation in pg/mL (2 experiments, n = 8 per group except naive, n = 4).
